# HSP110 dependent HSP70 disaggregation machinery mediates prion-like propagation of amyloidogenic proteins in metazoa

**DOI:** 10.1101/795435

**Authors:** Jessica Tittelmeier, Carl Alexander Sandhof, Heidrun Maja Ries, Silke Druffel-Augustin, Axel Mogk, Bernd Bukau, Carmen Nussbaum-Krammer

## Abstract

The gradual accumulation and prion-like propagation of α-synuclein and other amyloidogenic proteins is associated with devastating neurodegenerative diseases. The metazoan disaggregation machinery, a specific combination of HSP70 and its co-chaperones, is able to disassemble α-synuclein fibrils *in vitro*, but the physiological consequence *in vivo* is unknown. To explore this, we used *Caenorhabditis elegans* models that exhibit pathological features of α-synuclein, such as misfolding, intercellular spreading and toxicity. We inhibited the HSP70 disaggregase by depleting the crucial component HSP-110 and monitored the effect on α-synuclein related phenotypes. The knockdown of HSP-110 not only impaired HSP70 disaggregation activity and prevented the resolubilization of amorphous heat shock induced firefly luciferase aggregates, but also compromised the cellular folding capacity. In stark contrast, HSP-110 depletion reduced α-synuclein foci formation, cell-to-cell transmission and toxicity. Similar effects were observed for a polyQ model substrate, confirming that inhibition of HSP70 disaggregation function mitigates amyloid toxicity. These data demonstrate that the metazoan HSP70 disaggregation complex plays a critical role in the prion-like propagation of amyloid-type conformers. Therefore, the HSP70 disaggregation activity is a double-edged sword as it is essential for the maintenance of cellular proteostasis while being involved in the generation of toxic amyloid-type protein species.

## INTRODUCTION

Prion proteins can adopt alternative conformations with the ability to self-propagate. This concept of self-templated replication of misfolded protein species was first described for pathogenic protein aggregates in mammalian prion diseases and revolutionized our understanding of infectious agents (Griffith, 1967; Prusiner, 1982). Since their discovery in humans and animals, prions have been found in diverse other organisms, including yeast and bacteria (Wickner, 1994; Yuan and Hochschild, 2017). Work on yeast prions has been instrumental to elucidate basic mechanisms underlying protein-based inheritance (Serio et al., 2000; Sparrer et al., 2000; Wickner et al., 2013).

The prion state replicates via a mechanism described as nucleation dependent polymerization (Jarrett and Lansbury, 1993; Serio et al., 2000; Sparrer et al., 2000), where the rate-limiting step is the initial formation of a seeding-competent oligomeric nucleus consisting of misfolded conformers. This seed then templates the conversion and catalyzes the incorporation of soluble monomers, leading to growth of oligomers into larger highly-ordered beta-sheet rich fibrillar assemblies termed amyloids. The typical architecture of amyloids where the β-strands of individual molecules stack on top of each other is central for their self-templated growth. This structure allows the addition of monomers at the ends of the fibrils. Since this is an intrinsic property of any amyloid-type aggregate, the prion concept is expandable to other amyloid forming proteins. Accordingly, also amyloidogenic proteins associated with neurodegenerative diseases are propagating like prions, including Aβ and tau in Alzheimer disease (AD), α-synuclein (α-Syn) in Parkinson disease (PD), or proteins containing expanded polyglutamine stretches (polyQ) in Huntington’s disease (HD) and other polyglutamine diseases (Aguzzi and Rajendran, 2009; Cushman et al., 2010; Prusiner, 2012).

In addition to seeded polymerization, there is increasing evidence that fragmentation is essential for efficient prion amplification (Pezza and Serio, 2007), because it exposes more fibril ends at which new monomers are added (Collins et al., 2004; Jarrett and Lansbury, 1993). The importance of fragmentation is exemplified by *in vitro* techniques of prion amplification, such as real-time quaking induced conversion (RT-QuIC) (Atarashi et al., 2008) and protein misfolding cyclic amplification (PMCA) (Saborio et al., 2001), where fragmentation is achieved mechanically by rigorous shaking or sonication, respectively. Moreover, only mathematical models that take fibril breakage into account are able to accurately predict amyloid growth kinetics observed *in vitro* and *in vivo* (Knowles et al., 2009; Masel et al., 1999).

Evidence for an essential role of fragmentation in prion propagation *in vivo* comes from studies in yeast, where faithful propagation of the prion state is dependent on the activity of molecular chaperones, in particular the AAA^+^ disaggregase HSP104 (Chernoff et al., 1995; Tuite and Lindquist, 1996). Inactivation of HSP104 by guanidine hydrochloride (GdnHCl) or deletion of the HSP104 gene results in prion loss (Chernoff et al., 1995; Grimminger et al., 2004; Tuite and Lindquist, 1996). Importantly, HSP104 does not function in isolation, but cooperates with the HSP70 system to dissolve protein aggregates (Glover and Lindquist, 1998; Masison and Reidy, 2015; Mogk et al., 2018). Mechanistically, substrates are recognized by the HSP70-DNAJ (HSP40) chaperone pair and then handed over to HSP104, which extrudes single polypeptides from the aggregate by threading them through the central pore of the HSP104 hexamer (Lum et al., 2004). This ATP-dependent disaggregation activity leads to the formation of smaller prion fragments, so-called propagons, which have an increased replication capacity and are also more efficiently transferred to daughter cells (Chernoff et al., 1995; Mogk et al., 2018; Satpute-Krishnan et al., 2007; Tessarz et al., 2008; Tuite and Lindquist, 1996; Winkler et al., 2012). Fragmentation is therefore crucial not only for the amplification of prion conformers at the molecular level, but also at the cellular level. The life cycle of amyloid-based prions seems to involve continuous cycles of seeded assembly and fragmentation and only a particular equilibrium of association and dissociation kinetics allow propagation of the aggregated form.

While previous work revealed that metazoan cells exhibit disaggregation activity and also contain all co-factors required for cytosolic propagation and dissemination of an ectopically expressed amyloid-like yeast prion (Cohen et al., 2006; Hofmann and Vorberg, 2013; Hofmann et al., 2013; Krammer et al., 2009a; Krammer et al., 2009b), the nature of these co-factor(s) was still unclear, since direct homologs of HPS104 are absent from the metazoan cytosol. The recent discovery of a metazoan disaggregation machinery in which a particular HSP110-type nucleotide exchange factor (NEF) cooperates with the human HSP70-DNAJ system to fragment and depolymerize amyloid aggregates (Gao et al., 2015; Scior et al., 2018) has raised the question of whether this activity could prevent or promote the prion-like propagation of amyloid-type aggregates in metazoans and what physiological implications this would have. Since prion-like amplification and intercellular spreading of disease-linked proteins is implicated in the propagation of the pathology in neurodegenerative diseases such as AD and PD, co-factors involved in this process might be potential therapeutic targets.

In this study, we have investigated the role of the HSP70 metazoan disaggregation machinery in aggregation and toxicity of amyloid-type aggregates in *Caenorhabditis elegans* (*C. elegans*). Diminishing its activity by tissue-specific knockdown (KD) of HSP-110 impaired the resolubilization of amorphous aggregates such as heat shock (HS)-induced firefly luciferase aggregates on the one hand, while mitigating the formation of toxic spreading-competent α-Syn species on the other hand. These results reveal that a delicate balance between seeded polymerization and chaperone mediated fragmentation is necessary for efficient propagation of prion-like conformers in metazoan tissues and suggest that selectively interfering with chaperone-mediated remodeling of amyloid-type substrates might be beneficial because it interrupts the vicious circle of seeded polymerization and fragmentation.

## RESULTS

### Tissue-specific KD of HSP-110 prevents disassembly of HS-induced firefly luciferase aggregates

Our previous work revealed that a specific human HSP70-DNAJ-HSP110 network exhibits *in vitro* disaggregation activity towards amyloid α-Syn fibrils and detergent insoluble α-Syn extracted from *C. elegans* (Gao et al., 2015). However, its effect on aggregation and toxicity of α-Syn *in vivo* has yet to be addressed. Two scenarios are conceivable. The disaggregation activity could either lead to a resolubilization and refolding of aggregated α-Syn, which would result in reduced toxicity. Alternatively, it could contribute to the multiplication of α-Syn aggregates by continuously generating new seeds by fragmentation, analogous to the function of HSP104 and the HSP70-DNAJ system in yeast prion propagation, which would likely increase toxicity.

In an effort to assess the role of the metazoan disaggregation machinery in amyloid aggregate propagation *in vivo*, we employed *C. elegans* as model organism. Members of the nematode disaggregation system are highly conserved, but less redundant than in the human system (Kirstein et al., 2017). Yet, the cognate HSP70 of the cytosol and nucleus (HSP-1) is involved in a variety of different essential processes in addition to disaggregation, including protein synthesis, clathrin-mediated endocytosis, or chaperone-mediated autophagy (Stricher et al., 2013). The manipulation of this core component of the HSP70 network would lead to a plethora of side effects. Importantly, cooperation of the HSP70 with particular co-chaperones of the DNAJ-protein family (class B-type) and the HSP110-type family of NEFs determines functional specificity with respect to disaggregation (Gao et al., 2015; Mogk et al., 2018). While there are several DNAJ co-chaperones that are functionally redundant, *C. elegans* has only one cytosolic HSP110-type NEF (HSP-110), which is a key component of the HSP70 disaggregation machinery (Gao et al., 2015; Kirstein et al., 2017; Mogk et al., 2018; Rampelt et al., 2012). By manipulating HSP-110 levels, the disaggregation activity of the HSP70 chaperone network can be specifically modulated. However, since an *hsp-110* knockout is lethal and a systemic knockdown (KD) by RNA interference (RNAi) affects growth, development, and fertility (Kamath et al., 2003), we decided to restrict the KD of HSP-110 to the same tissue that expresses the selected substrate protein (e.g. α-Syn). This allows us both to focus on the specific effect on the co-expressed protein of interest and to reduce unwanted side effects in other tissues. To this end, we generated *C. elegans* strains expressing an RNAi hairpin (HP) construct targeting the *hsp-110* mRNA specifically in muscle cells (HPI and HPII) (Winston et al., 2002). To ensure tissue-specific KD of *hsp-110* and to prevent systemic RNAi, all strains harbor a loss-of-function mutation in *sid-1*, a gene coding for a double stranded RNA transporter, which is essential for inter-tissue transport of dsRNA and systemic RNAi (Winston et al., 2002). To verify a tissue-specific KD of *hsp-110*, we introduced both *hsp-110* hairpin constructs into a strain with endogenously GFP-tagged HSP-110 (HSP-110::GFP). Expression of both *hsp-110* hairpin constructs resulted in the loss of GFP fluorescence exclusively in muscle cells, confirming a tissue-specific KD (**Fig. 1A**).

**Figure 1.**
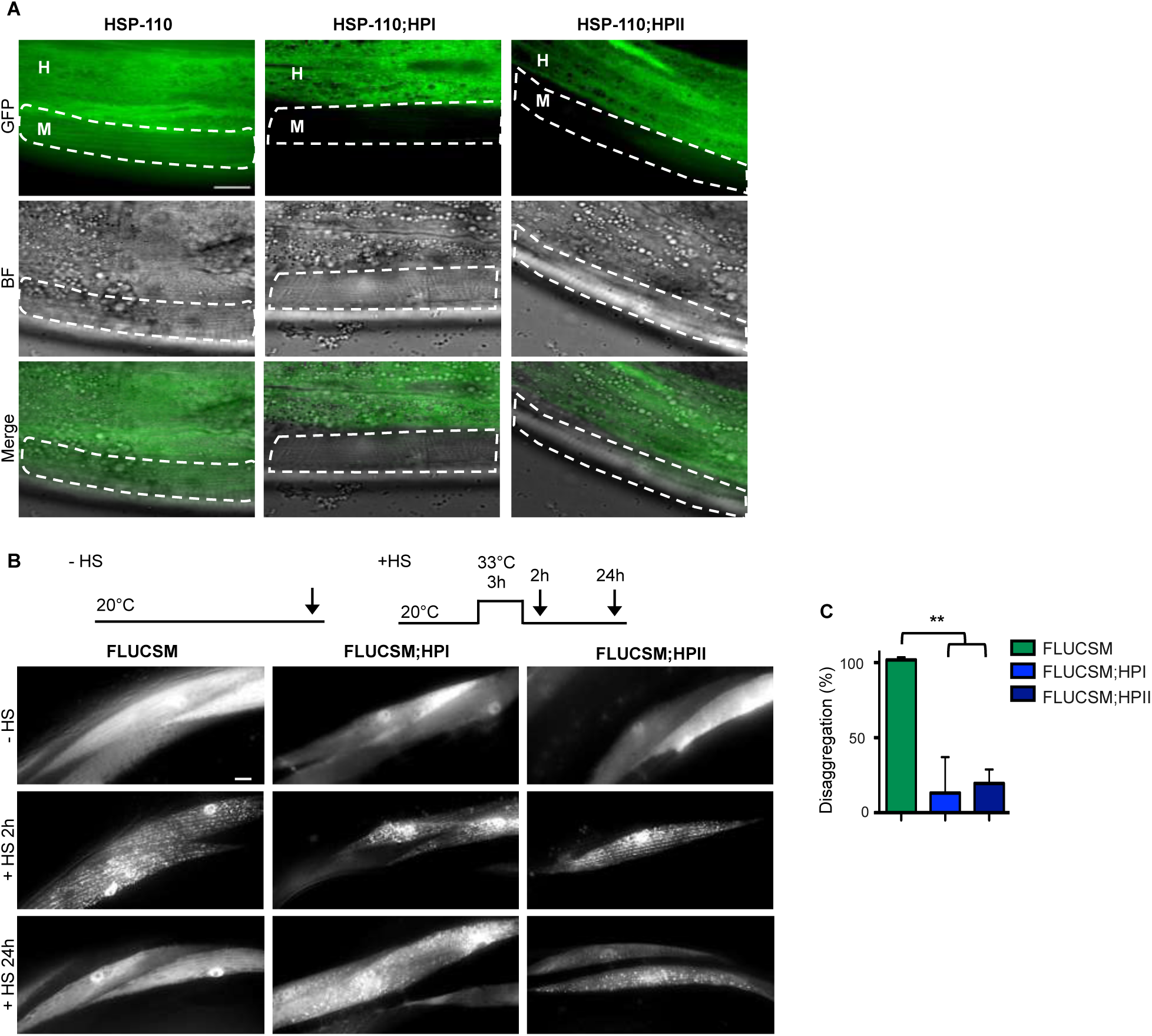
Tissue-specific KD of HSP-110 prevents resolubilization of HS-induced firefly luciferase aggregates. **A** Single plane confocal fluorescent microscopy images of 5-day-old animals harboring GFP-tagged endogenous HSP-110 (HSP-110::GFP). Muscle cells are outlined. M: muscle, H: hypodermis. In animals that co-express hairpin constructs targeting *hsp-110* in muscle cells (HSP-110;HPI and HSP-110;HPII), GFP fluorescence is strongly reduced in muscle cells in comparison to control animals, indicating HSP-110::GFP depletion specifically in muscle cells. Scale bar: 10 µm. **B** Experimental set-up: Age-synchronized 4-day-old animals were subjected to heat stress for 3 h at 33°C (+HS) then returned to 20°C or left at 20°C (-HS), respectively. Arrows indicate imaging time points. The -HS controls were imaged at the same time point as the +HS animals after 24h at 20°C. Maximum intensity projections of fluorescent microscopy z-stacks of animals expressing the indicated transgenes. HS-induced FLUCSM::GFP foci are cleared after a recovery for 24h at 20°C in the WT background, but persist in the presence of the *hsp-110* hairpin construct. Scale bar: 20 µm. **C** Quantification of FLUCSM::GFP foci disaggregation. % disaggregation is calculated as 100 - ratio of FLUCSM::GFP foci area relative to total muscle area at the +HS 24h time point to the +HS 2h time point. Data are displayed as mean ± SEM (in %). Statistical analysis was done using one-way ANOVA with Dunnett’s adjustment for multiple comparisons. ** p ≤ 0.01. All strains harbor the *sid-1(pk3321)* allele.

We then examined whether HSP-110 KD impairs the cellular disaggregation activity by using a well characterized firefly luciferase reporter (FLUCSM::EGFP) expressed in muscle cells as a model substrate (Gupta et al., 2011). FLUCSM::EGFP remained soluble under ambient growth conditions, but formed numerous aggregates upon acute heat shock (3h 33°C) in both wild-type (WT) and *hsp-110* hairpin expressing animals (**Fig. 1B**). After a 24 hour recovery period, FLUCSM::EGFP foci were almost completely dissolved in the WT background, whereas a large fraction of muscle cells still harbored multiple FLUCSM::EGFP foci upon expression of *hsp-110* hairpins (**Fig. 1B and C**). Thus, the HSP-110 KD led to persistence of heat-induced amorphous FLUCSM aggregates, indicating that disaggregation activity was indeed impaired (Rampelt et al., 2012).

### KD of HSP-110 reduces cytosolic α-Syn and Q35 foci formation and toxicity

We next tested the effect of impaired disaggregation activity on an amyloidogenic substrate. After introducing the *hsp-110* hairpin constructs into animals expressing yellow fluorescent protein (YFP)-tagged α-Syn (α-Syn::YFP) in muscle cells (van Ham et al., 2008), we analyzed α-Syn::YFP aggregation and toxicity at day 4, 5 and 6 of life (corresponding to day 1, 2 and 3 of adulthood). In line with previous reports (van Ham et al., 2008), α-Syn::YFP accumulated into large distinct foci (**Fig. 2A, 2B, S1A** and **S1B**), which was associated with a progressive decline of muscle function as assessed by a motility assay (**Fig. 2C** and **S1C**). Remarkably, the co-expression of *hsp-110* hairpins significantly reduced the number of foci and rescued the toxicity of α-Syn::YFP (**Fig. 2A, 2B** and **2C**). This suggests that HSP-110-dependent disaggregation activity is required for α-Syn::YFP foci formation and toxicity.

**Figure 2.**
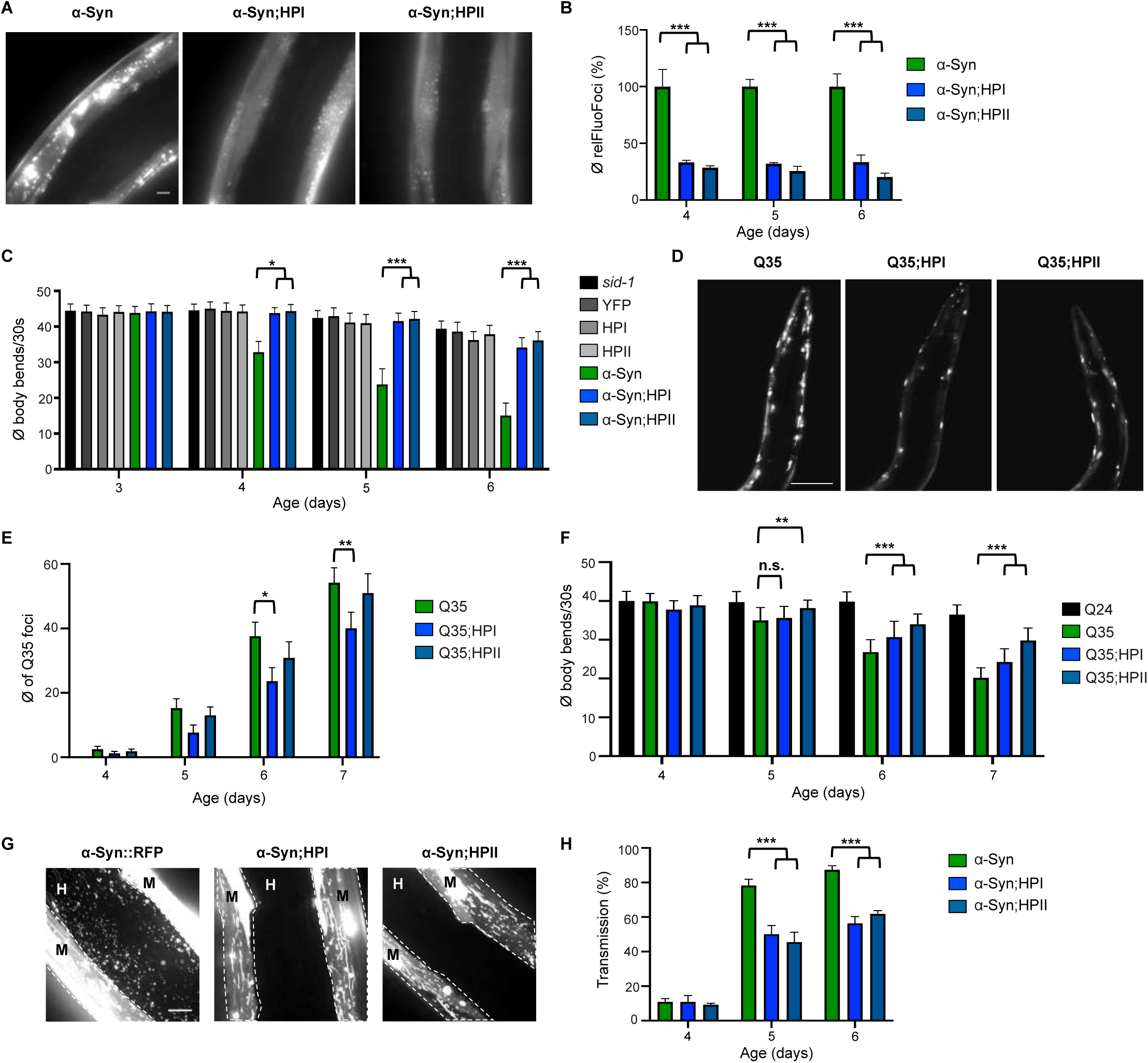
KD of HSP-110 reduces cytosolic α-Syn and Q35 foci formation and toxicity. **A** Maximum intensity projections of fluorescent microscopy z-stacks of 5-day-old nematodes expressing the indicated transgenes are shown. Scale bar: 10 µm. The co-expression of HPI and HPII reduces the amount of α-Syn::YFP inclusions. **B** Quantification of α-Syn::YFP foci in 4-, 5- and 6-day-old animals. Product of mean foci fluorescence and foci area relative to muscle area is displayed (relFluoFoci). Data are shown as mean ± SEM (in %). Statistical analysis was done using two-way ANOVA with Tukey’s multiple comparison test. *** p ≤ 0.001. **C** Motility Assay as a measure for transgene toxicity. Displayed is the mean number of body bends per 30 s ± SEM. Statistical analysis was done using two-way ANOVA with Tukey’s multiple comparison test. * p ≤ 0.05, *** p ≤ 0.001. **D** Maximum intensity projections of fluorescent microscopy z-stacks of 6-day-old animals expressing the indicated transgenes. Scale bar: 100 µM. The co-expression of HPI and HPII reduces the amount of Q35::YFP inclusions. **E** Quantification of Q35::YFP foci in WT and *hsp-110* hairpin background. Shown is the mean ± SEM. Statistical analysis was done using two-way ANOVA with Dunnett’s multiple comparison test. * p ≤ 0.05, ** p ≤ 0.01. **F** Motility Assay as a measure for transgene toxicity. Displayed is the mean number of body bends per 30 s ± SEM. Statistical analysis was done using one-way ANOVA with Tukey’s multiple comparisons test. n.s. = not significant, ** p ≤ 0.01, *** p ≤ 0.001. **G** Maximum intensity projections of fluorescent microscopy z-stacks of of 5-day-old nematodes expressing the indicated transgenes. White dashed lines outline the borders of muscle cells. Scale bar: 10 µm. M: muscle, H: hypodermis. Signal outside of muscle cells reveals spreading of α-Syn. **H** Quantification of animals showing α-Syn transmission at indicated ages. Displayed is the mean ± SEM (in %). Statistical analysis was done using two-way ANOVA with Tukey’s multiple comparison test. *** p ≤ 0.001. All strains harbor the *sid-1(pk3321)* allele, except for the YFP only and Q24 controls.

We next asked whether this effect was specific to α-Syn or whether the same could be observed with another amyloidogenic protein. To test this, we introduced the *hsp-110* hairpins into animals expressing 35 polyglutamine (polyQ) residues fused to YFP (Q35::YFP), another substrate that forms fibrillar aggregates in an age-dependent manner (Morley et al., 2002). KD of HSP-110 also reduced the number of foci and toxicity of Q35 (**Fig. 2D, 2E, 2F, S1D, S1E** and **S1F**). In sum, HSP-110 activity appears to enhance α-Syn and Q35 foci formation and toxicity.

Protein quality control pathways are typically intertwined and inhibiting one pathway sometimes results in upregulation of other components, which could indirectly prevent α-Syn and Q35 misfolding and toxicity in the *hsp-110* KD strains. However, comparable expression levels of typical stress inducible chaperone genes verified that the KD of HSP-110 did not activate the stress transcription factors HSF-1/HSF1 or DAF-16/FOXO (**Fig. S2**). Since total protein levels of α-Syn::YFP and Q35::YFP remained largely unchanged (**Fig. S3**), it could also be ruled out that an enhanced degradation was responsible for the observed reduction of foci formation and toxicity. Together with the observations on luciferase reactivation, these data indicate that the effect of HSP-110 KD was specific to its role in substrate disaggregation and was not mediated by indirect compensatory mechanisms.

HSP104-dependent remodelling of yeast prion aggregates also promotes efficient segregation of seeds into daughter cells, ensuring inheritance of the prion state (Satpute-Krishnan et al., 2007). To test whether the inhibition of HSP-110 dependent disaggregation equally affects α-Syn dissemination in metazoans, we employed our recently developed α-Syn spreading model to monitor the transmission of α-Syn from donor muscle cells into recipient hypodermal cells (Sandhof et al., 2019). The dissemination of α-Syn is age-dependent and can be detected in less than 10% of the animals on day 4 and in more than 90% on day 5 and later in life (Sandhof et al., 2019). KD of HSP-110 significantly reduced the transmission of α-Syn particles between tissues with protein transfer being detected in only 50% of *hsp-110* hairpin animals on day 5 and 6 compared to up to 90% of WT animals (**Fig. 2G** and **2H**). Hence, in the absence of HSP-110, α-Syn is inefficiently transferred to neighbouring tissues. Since polyQ proteins are not spreading to neighbouring tissues in our model system (Sandhof et al., 2019), we could not assess the effect of HSP-110 levels on Q35 spreading. Nevertheless, these findings revealed that the HSP-110 dependent disaggregation activity seems to remodel α-Syn aggregates to create toxic species that can transfer between cells.

Overall, these results support the notion that the KD of HSP-110 impairs the HSP70 disaggregation machinery, which on the one hand prevents the removal of amorphous aggregates, and on the other hand abolishes the propagation and intercellular transmission of toxic amyloid-type substrates.

### WT and ATPase deficient yeast SSE1 can compensate the loss of *C. elegans* HSP-110

What is the specific contribution of HSP-110 to the observed disaggregation activity in *C. elegans*? HSP110s have a substrate binding and a nucleotide binding domain similar to HSP70, therefore an ATP-driven chaperone-like activity has been proposed for this chaperone family (Mattoo et al., 2013; Shorter, 2011). Indeed, structurally unrelated NEFs, such as Snl1ΔN (BAG-1 homolog) or FES-1 (HSPBP1 homolog) were not able to efficiently substitute an HSP110-family NEF in disaggregation (Kaimal et al., 2017; Rampelt et al., 2012). However, previous results revealed that only the nucleotide exchange activity is necessary for efficient substrate disaggregation *in vitro*, while its intrinsic ATPase activity is dispensable (Rampelt et al., 2012). To investigate whether this also applies to *C. elegans*, we took advantage of the high conservation of the chaperones involved in disaggregation, which allows individual members to cooperate across species boundaries (Rampelt et al., 2012). Since yeast NEFs were shown to be able to replace the human HSP110-type NEF *in vitro* (Rampelt et al., 2012), we introduced an extrachromosomal array coding for yeast WT or ATPase deficient K69M mutant SSE1 (HSP110-type NEF) or FES1 (armadillo-type NEF) into α-Syn::YFP and *hsp-110* hairpin expressing animals by microinjection. We found that WT and K69M mutant SSE1 could complement for the KD of the endogenous *C. elegans* HSP-110 and restore the formation of visible α-Syn foci (**Fig. 3A**) and α-Syn toxicity (**Fig. 3B**). Since the transgenes were expressed from an extrachromosomal array, which results in variable expression levels in a variable number of cells, a full complementation was not expected. Expression of FES1 on the other hand had no obvious impact on α-Syn foci formation (**Fig. 3A**) and did not abrogate the suppression of α-Syn toxicity in the HSP-110 KD background (**Fig. 3B**). Only an HSP110-type NEF seems to be able to reestablish the α-Syn phenotype as in WT, however, we cannot rule out that yeast FES1 is not able to cooperate with the *C. elegans* HSP70 system. Therefore, we subsequently focused only on the SSE1 variants and tested their impact on α-Syn spreading in our α-Syn::RFP model. With the introduction of WT and K69M mutant SSE1 into the *hsp-110* hairpin animals the intercellular transmission of α-Syn was also restored (**Fig. 3C**).

**Figure 3.**
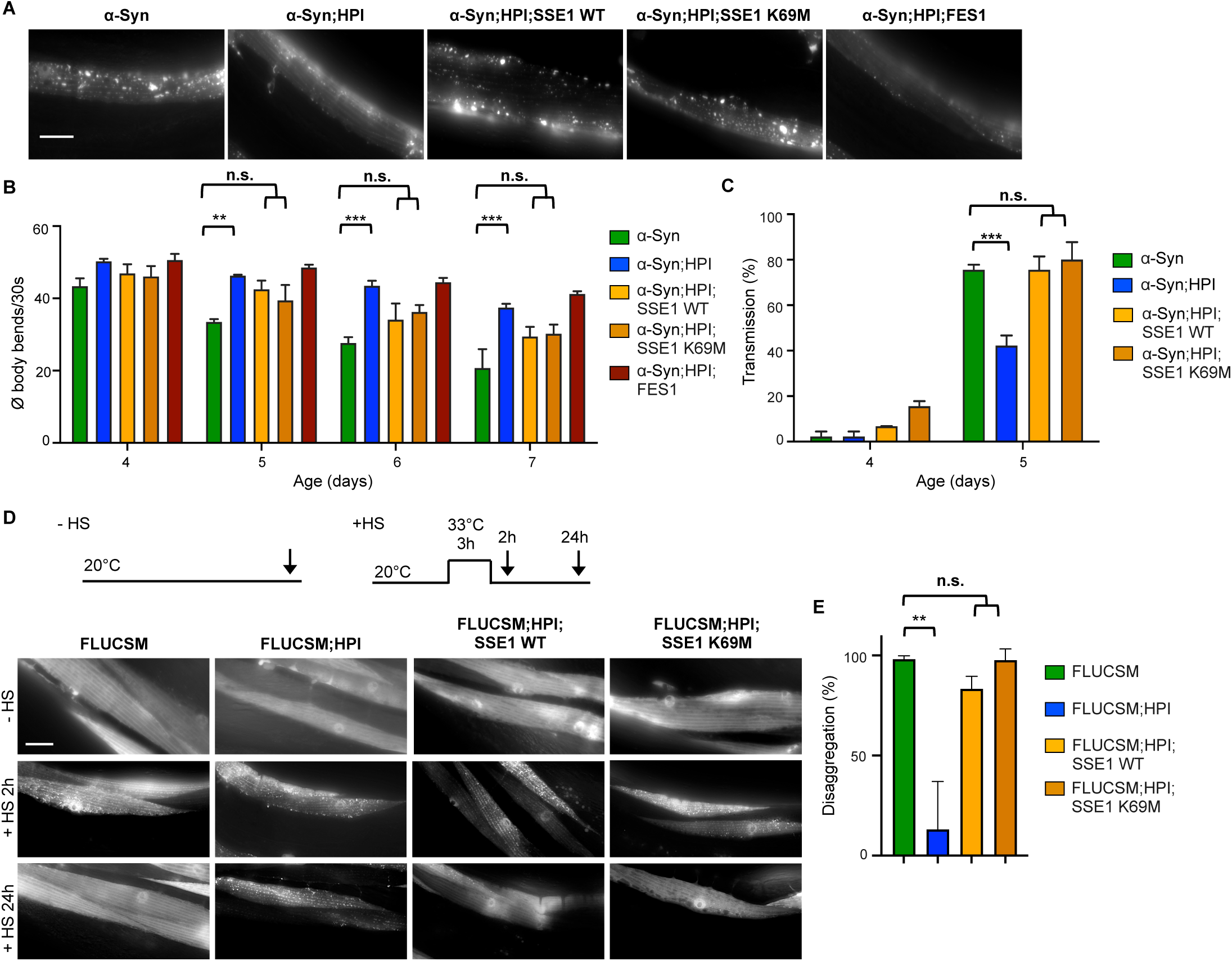
WT and ATPase deficient yeast SSE1 can compensate the loss of *C. elegans* HSP-110. **A** Maximum intensity projections of confocal microscopy z-stacks of 5-day-old animals expressing the indicated transgenes. Scale bar: 10 µm. The expression of yeast WT and K69M mutant SSE1 but not FES1 reversed the reduction of α-Syn::YFP foci in the *hsp-110* hairpin expressing animals. **B** Quantification of motility as a measure for transgene toxicity. Displayed is the mean ± SEM amount of body bends/30s of animals expressing the indicated transgenes. Statistical analysis was done using two-way ANOVA and with Bonferroni’s multiple comparison test. n.s. = not significant, ** p ≤ 0.01, *** p ≤ 0.001. The presence of yeast WT and K69M mutant SSE1 but not FES1 suppressed the rescue of α-Syn::YFP toxicity in *hsp-110* hairpin expressing animals. **C** Quantification of animals showing α-Syn transmission into the hypodermis at indicated ages. Displayed is the mean ± SEM (in %). Statistical analysis was done using two-way ANOVA with Bonferroni’s multiple comparison test. n.s. = not significant, *** p ≤ 0.001. **D** Experimental set-up: Age-synchronized 4-day-old animals were subjected to heat stress for 3 h at 33°C (+HS) then returned to 20°C or left at 20°C (-HS), respectively. Arrows indicate imaging time points. The -HS controls were imaged at the same time point as the +HS animals after 24h at 20°C. Maximum intensity projections of fluorescent microscopy z-stacks of animals expressing the indicated transgenes. Scale bar: 20 µm. The impaired clearance of HS-induced FLUCSM::GFP foci in the *hsp-110* hairpin expressing cells is rescued by co-expression of WT and K69M mutant SSE1. **E** Quantification of FLUCSM::GFP foci disaggregation. % disaggregation is calculated as 100 - ratio of FLUCSM::GFP foci area relative to total muscle area at the +HS 24h time point to the +HS 2h time point. Data are displayed as mean ± SEM (in %). Statistical analysis was done using one-way ANOVA with Dunnett’s multiple comparison test. n.s. = not significant, ** p ≤ 0.01. All strains harbor the *sid-1(pk3321)* allele.

To verify whether the observed effects with the SSE1 variants could be attributed to a reestablishment of disaggregation activity, the FLUCSM::EGFP reporter was used as an amorphous substrate. As before, FLUCSM::EGFP remained soluble in all strains under non-HS conditions, but readily formed multiple distinct foci after being subjected to heat stress (**Fig. 3D** and **3E**). Again, HSP-110 KD animals were not able to resolubilize aggregated FLUCSM::EGFP during the 24h recovery period and numerous aggregates could still be observed in each cell (**Fig. 3D** and **3E**). Additional expression of either WT or ATPase dead SSE1 led to the removal of almost all visible foci during the recovery period and muscle cells showed diffuse FLUCSM::EGFP fluorescence with only sporadic foci in a small number of cells (**Fig. 3D** and **3E**).

Taken together, these observations show that the yeast HSP-110 homolog SSE1 can cooperate with the *C. elegans* Hsp70-DNAJ system *in vivo* and compensate the loss of endogenous HSP-110, leading to an increase of α-Syn foci formation, spreading and toxicity as well as the solubilization of HS-induced FLUCSM::EGFP foci. Moreover, in line with previous *in vitro* data, the sole NEF activity appears to be sufficient to promote disaggregation whereas the intrinsic ATPase activity of HSP110 is likely to be of no importance for its role in aggregate disaggregation (Rampelt et al., 2012). This argues against a specific chaperone-like contribution of HSP-110 to disaggregation.

### KD of HSP-110 impairs cellular protein folding homeostasis

We further investigated whether the KD of HSP-110 might cause more general cellular defects that impair functionality of the affected tissue. The loss of HSP-110 could negatively affect cellular HSP70 capacity, as HSP70 is not efficiently released from its substrates in the absence of a NEF. This would reduce the pool of HSP70 available for its other functions and therefore, as a negative side effect, KD of HSP-110 might interfere with the cellular folding environment. To test this, the *hsp-110* hairpins were introduced into strains each containing an endogenous protein variant with a temperature sensitive (ts) mutation leading to folding defects (Ben-Zvi et al., 2009; Gidalevitz et al., 2006). Strains harboring either paramyosin(ts) [*unc-15*(*e1402*)], myosin(ts) [*unc-54*(*e1301*)], or perlecan(ts) [*unc-52*(*e669*, *su250*)], have been established as sensors that provide information about the folding capacity of the proteostasis network (Ben-Zvi et al., 2009; Gidalevitz et al., 2006). These animals behave as superficially WT at permissive temperatures (15°C), but exhibit severe movement defects due to disrupted muscle filaments resulting from the aggregation of the ts mutant proteins at restrictive temperatures (25°C) or under conditions that impair the cellular folding environment (Ben-Zvi et al., 2009; Gidalevitz et al., 2006).

The respective ts phenotypes were confirmed at 25°C, while the control and *hsp-110* hairpin animals with WT *unc* alleles showed no abnormalities (**Fig. 4A, 4B, S4** and **S5**). At the permissive temperature of 15°C, control, *hsp-110* hairpin and ts mutant animals were almost indistinguishable and did not exhibit any obvious phenotypes (**Fig. 4A, 4B, S4** and **S5**). In contrast, movement defects were readily evident in ts mutant strains with reduced HSP-110 levels (**Fig. 4B, S4B** and **S4C**). The decline in motility in the latter animals correlated with distorted muscle filaments caused by the misfolding and misassembly of the respective ts mutant proteins (**Fig. 4A** and **S4A**). Thus, endogenous metastable protein variants, which are continuously misfolding even at low ambient growth temperatures, are no longer correctly (re-)folded upon HSP-110 KD, leading to exposure of the ts mutant phenotype at permissive temperatures.

**Figure 4.**
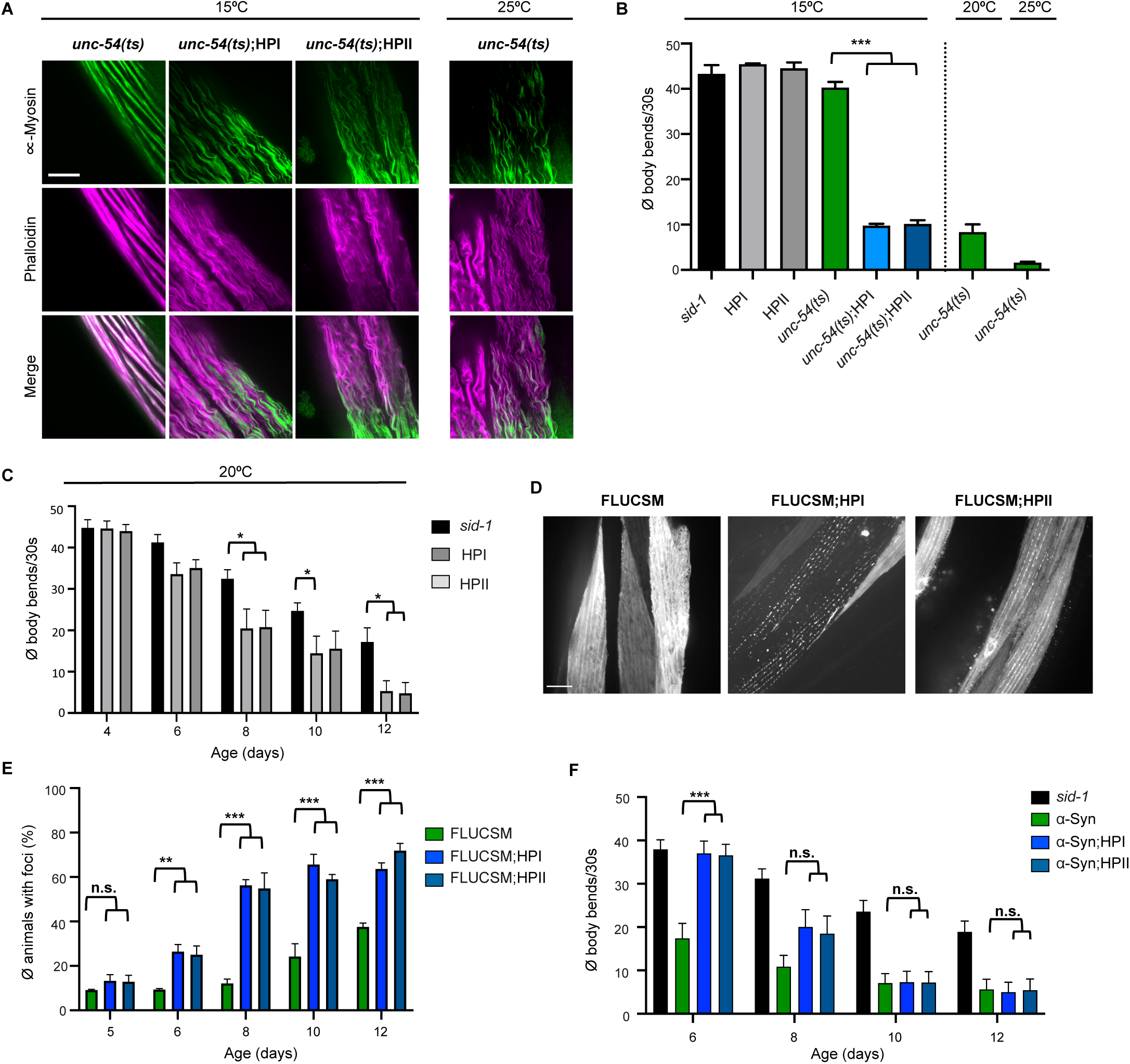
HSP-110 HP expression impairs cellular protein folding homeostasis. **A** Maximum intensity projections of confocal z-stacks of 4-day-old animals expressing the indicated transgenes or harboring the indicated temperature-sensitive (ts) mutations. Muscle cells were stained with anti-myosin (green) and Alexa Flour 647-phalloidin (purple). Scale bar: 5 µm. Myosin misfolding reflects the exposure of the ts mutant phenotype of myosin(ts) [*unc-54*(*e1301*)] at the permissive temperate of 15°C by co-expression of the *hsp-110* hairpins. **B** Quantification of motility as a measure for toxicity. Displayed is the mean ± SEM of body bends/30s of animals expressing the indicated transgenes or harboring the indicated mutations. Statistical analysis was done using one-way ANOVA with Dunnett’s multiple comparison test. *** p ≤ 0.001. Co-expression of the *hsp-110* hairpins exposed the ts mutant phenotype at the permissive temperate of 15°C, resulting in a significant increase in movement defects. All strains contain the *sid-1(pk3321)* allele. **C** Quantification of motility as a measure for transgene toxicity. Displayed is the mean ± SEM amount of body bends/30s of animals expressing the indicated transgenes or harboring the indicated mutations. Statistical analysis was done using two-way ANOVA with Tukey’s multiple comparison test. * p ≤ 0.05. Expression of the *hsp-110* hairpins caused movement defects with increasing age. **D** Maximum intensity projections of confocal z-stacks of 12-day-old animals expressing the indicated transgenes. Scale bar: 10 µm. FLUCSM forms foci with increasing age, which is markedly accelerated upon *hsp-110* KD. **E** Quantification of animals with FLUCSM::GFP foci at indicated days of life. Data are displayed as mean ± SEM (in %). Statistical analysis was done using two-way ANOVA with Dunnett’s multiple comparison test. n.s. = not significant, ** p ≤ 0.01, *** p ≤ 0.001. **F** Quantification of motility as a measure for transgene toxicity. Displayed is the mean ± SEM amount of body bends/30s of animals expressing the indicated transgenes or harboring the indicated mutations. Statistical analysis was done using two-way ANOVA with Tukey’s multiple comparison test. n.s. = not significant, *** p ≤ 0.001. Expression of the *hsp-110* hairpins caused movement defects with increasing age. The beneficial effects of HSP-110 KD on amyloid toxicity is masked in older animals. All strains harbor the *sid-1(pk3321)* allele.

These results suggest that the KD of HSP-110 impairs not only HSP70 disaggregation activity but also HSP70 (re-)folding activity, which negatively impacts the cellular folding environment.

### KD of HSP-110 impairs muscle function with increasing age

We asked whether an impairment of the disaggregation machinery affects muscle cell function during aging since there is stronger demand on the proteostasis network with the cumulative amount of misfolding proteins (David et al., 2010; Walther et al., 2015). To investigate muscle function of animals with a compromised disaggregation machinery, animals were subjected to a motility assay under both normal (20°C) and elevated ambient temperature (25°C) to slightly challenge the proteostasis network. At both 20°C and 25°C growth conditions, there is no difference between control and *hsp-110* hairpin strains in young animals (**Fig. 2C, 4C, S5A** and **S6A**). However, during aging there is a faster decline in motility in *hsp-110* hairpin animals compared to the control strains (**Fig. 4C** and **S6A**). This is likely due to an accelerated collapse of proteostasis when the disaggregation system is compromised. Accordingly, also the age-dependent FLUCSM foci formation is accelerated in the *hsp-110* KD background (**Fig. 4D** and **4E**). As a result, the rescue of α-Syn toxicity by the KD of HSP-110 observed in younger animals is lost in older animals (**Fig. 4F**). This cannot be attributed to a restoration of disaggregation activity and an increase in α-Syn foci in older animals, since the levels of HSP-110 and α-Syn foci are still significantly reduced in the *hsp-110* hairpin strains (**Fig. S6B, S6C,** and **S6D**).

These data suggest that in young animals, the low level of misfolded endogenous proteins does not overwhelm cellular protein folding capacity where other members of the proteostasis network can still buffer the impaired HSP-110 activity. At this age, only the expression of amyloidogenic proteins pose a significant challenge for the proteostasis network since the protective effect of KD of HSP-110 in α-Syn and Q35 transgenic animals is revealed despite impairment of the essential disaggregation activity. In contrast, in older, post-reproductive, animals the increasing load of endogenous misfolded proteins represents an additional burden on the disaggregation system that eventually masks the positive effects on α-Syn and Q35 toxicity. These results illustrate that the phenotype resulting from the impairment of the disaggregation system depends on the nature of the predominant substrate(s) at a given time.

## DISCUSSION

There is growing evidence that many amyloidogenic aggregates associated with prevalent neurodegenerative diseases such as AD and PD can propagate like prions and spread from cell to cell, which might accelerate disease progression. However, the mechanisms that drive these processes are still largely unknown. The yeast prion model system paved the way to identify and understand cellular factors of prion propagation, leading to the seminal discovery that efficient prion propagation is inevitably linked to amyloid fragmentation by the AAA^+^ disaggregase HSP104 (Chernoff et al., 1995). It remained a major open question whether similarly acting cellular chaperones exist in metazoa, which lack HSP104.

The present study provides proof-of-principle that the HSP70 disaggregation machinery is required for the prion-like propagation of disease-associated amyloidogenic proteins in metazoa. Blocking the disaggregation activity by knocking down HSP-110 interfered with the formation of toxic α-Syn and Q35 conformers in *C. elegans* (**Fig. 5A**). The visible cytosolic foci in WT animals likely constitute a reservoir, from which propagons are continuously released by the HSP70 disaggregase (**Fig. 5B**). KD of HSP70 also reduced α-Syn spreading in *C. elegans*, indicating that particles generated by disaggregation are preferred substrates for intercellular transfer. Indeed, smaller subfibrillar prion species are most infectious and more efficiently transported between cells than larger assemblies (Bett et al., 2017; Silveira et al., 2005).

**Figure 5.**
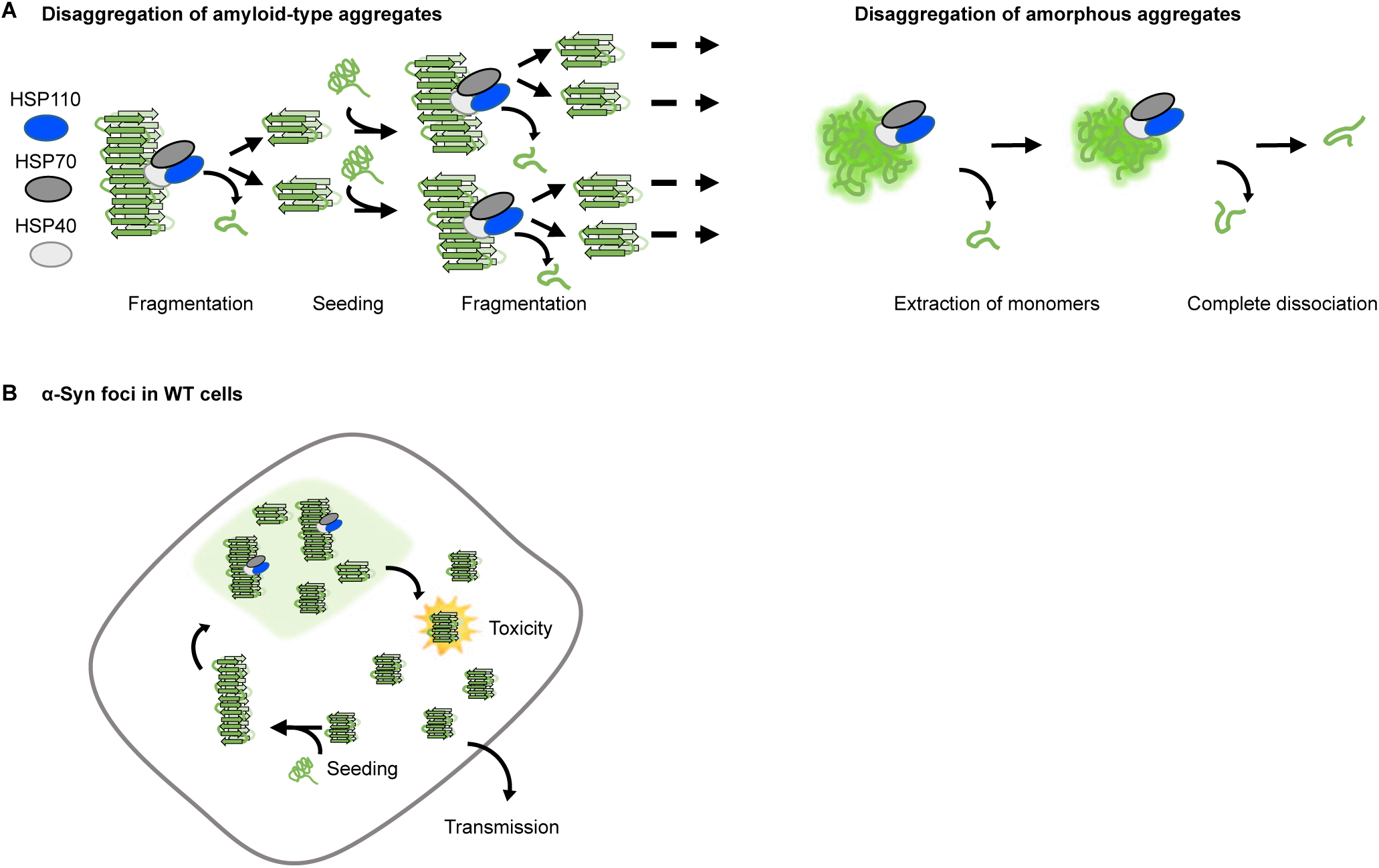
Model of the effect of disaggregation activity on aggregation and toxicity of amyloid-like and amorphous aggregates. **A** The disaggregation of amyloid-like aggregates by the HSP70 system is an important part of a propagation cycle involving repeated fragmentation of larger fibrils to generate and amplify toxic seeding-competent particles. In contrast, the disaggregation of amorphous aggregates leads to the formation of smaller seeding-incompetent fragments, which are eventually completely dissociated. **B** α-Syn foci in WT cells might constitute active sites for propagon formation. Disaggregation in these sites generates smaller α-Syn fragments that are more toxic and more readily transferred to adjacent tissues. As they are also more seeding competent, this leads to an amplification of the aggregated material.

The KD of HSP-110 in *C. elegans* resembles in several aspects an impairment of HSP104 in yeast. For example, HSP-110 depletion in *C. elegans* resulted in the persistence of heat-induced luciferase aggregates. Likewise, inhibition of HSP104 activity by GdnHCl also suppresses resolubilization of aggregated luciferase in heat shocked yeast cells (Ness et al., 2002). Impairing HSP-110 activity in *C. elegans* ameliorated the aggregation and toxicity of α-Syn and Q35. Similarly, the inhibition of HSP104 reduces aggregation and toxicity of other disease-associated amyloidogenic proteins, such as polyQ in yeast (Krobitsch and Lindquist, 2000). In addition, the HSP-110 dependent disaggregation activity seems to affect α-Syn aggregates similarly to how HSP104 remodels yeast prion complexes, leading to the amplification of (toxic) conformers that have an increased potential to transfer to neighbouring or daughter cells (Chernoff et al., 1995; Tuite and Lindquist, 1996). Since the HSP104-HSP70 system is known to fragment prion aggregates, it is tempting to speculate that the HSP70 disaggregase also fragments amyloidogenic substrates in *C. elegans*. Although HSP104 was the first chaperone discovered to be essential for prion propagation (Chernoff et al., 1995; Tuite and Lindquist, 1996), it does not work in isolation, but collaborates with the HSP70 system to disaggregate both amyloid and amorphous aggregates (Masison and Reidy, 2015; Winkler et al., 2012). Therefore, HSP70 co-chaperones also play a role in prion propagation and amyloid toxicity, including the yeast HSP110 SSE1 (Du et al., 2017; Fan et al., 2007; Kaimal et al., 2017; Kryndushkin and Wickner, 2007; Sadlish et al., 2008; Sun et al., 2011).

Recent work showed that the overexpression of an HSP110-type NEF extended the survival of mice expressing G85R mutant superoxide dismutase 1 (SOD1), which is linked to amyotrophic lateral sclerosis (ALS) (Nagy et al., 2016). The authors concluded that this is likely boosting the function of the disaggregation machinery, but without providing evidence for an increased activity. These results seem to contradict the present study at first, as SOD1 is a presumably amyloidogenic protein such as α-Syn. However, our previous work *in vitro* (Rampelt et al., 2012), as well as observations of others *in vivo* (Kampinga and Craig, 2010; Nollen et al., 2000) imply that not only too low but also too high HSP-110 levels poison the HSP70 disaggregation reaction. This can be explained by too slow and too fast nucleotide release from HSP70, hampering the HSP70 ATPase cycle. Therefore, HSP-110 overexpression might rather interfere with disaggregation activity, leading to less SOD1 aggregation and toxicity according to our model.

Given the opposite effects on amorphous vs. amyloid substrates, it is not surprising that KD of HSP-110 can lead to contradictory results, even in the context of the same model system. HSP-110 has been previously identified in a genome-wide screen for proteostasis regulators in *C. elegans*, in which an organism-wide KD of HSP-110 reduced Q35 aggregation (Silva et al., 2011). In contrast, the same systemic KD resulted in increased Q35 aggregation in another study (Scior et al., 2018). The respective outcome is likely to depend on the timing and strength of the KD, and to what extent it affects the folding of proteins other than Q35, both in the same tissue and in neighboring tissues, which may also influence muscle cell proteostasis through non-autonomous mechanisms (Morimoto, 2019; Shai et al., 2014). Our results clearly show that the impairment of the disaggregation machinery rescues the aggregation and toxicity of amyloid-like substrates, but at the same time negatively impacts the removal of amorphous aggregates or folding of metastable proteins. These side effects may ultimately override the positive effect on amyloidogenic proteins, especially during aging.

Although chaperones are generally beneficial and indispensable, the ability of prions to exploit the disaggregation machinery in order to propagate in cells ranging from yeast to metazoan as shown here, renders this activity a double-edged sword in disease and aging (Jones and Tuite, 2005; True, 2006; Wentink et al., 2019). Overall, our results show that caution is advised when attempting to influence disaggregation activity for disease intervention, as boosting the activity could produce more toxic seeds, while inhibiting it could adversely affect the removal of amorphous aggregates and cell function. The higher redundancy of HSP110-type co-chaperones with three isoforms (HSP105 (HSPH1), APG2 (HSPH2), and APG1 (HSPH3)) in humans could be advantageous here, as only one member could be targeted, which would likely lead to fewer side effects, but could still be sufficient to shift the balance of amyloid propagation towards resolubilization or degradation. Another promising approach could be to specifically abolish the disaggregation of amyloids while leaving the (re-)folding and disaggregation of other substrates unaffected, which may be achievable by targeting substrate-specific Hsp70 co-chaperones. Furthermore, the tissue-specific promoter used in our model system mediated an early and continuous KD of HSP-110 parallel to substrate expression. Transient inhibition of disaggregation activity by a temporary drug administration could reduce amyloid aggregate load to levels that can be managed by cells while minimizing negative side effects. Therefore, the development of small molecules for pharmacological modulation of specific components of the disaggregation machinery could be a promising way to combat diseases caused by amyloidogenic proteins.

## ACKNOWLEDGEMENTS

We thank Cindy Voisine, Sabine Gilch, Hermann Schätzl and all members of the Nussbaum lab for their helpful discussion and constructive comments on the manuscript. This study is part of the PROTEST-70 project within the EU Joint Programme - Neurodegenerative Disease Research (JPND) project. This project is supported through the following funding organisations under the aegis of JPND - www.jpnd.eu: Germany, Bundesministerium für Bildung und Forschung (BMBF, 01ED1807A to B.B. and 01ED1807B to C.N-K.). This work was also funded by the Deutsche Forschungsgemeinschaft (DFG, SFB1036 TP20 to C.N.-K. and TP08 to A.M. and B.B.) and the AMPro program of the Helmholtz Gemeinschaft (to B.B.).

## MATERIALS AND METHODS

### Maintenance of *C. elegans* and age-synchronization

All animals were cultured using standard methods (Brenner, 1974). If not otherwise indicated, worms were grown on nematode growth medium (NGM) plates seeded with *E. coli* strain OP50 at 20°C. Animals were age-synchronized by bleaching or egg laying (Sandhof et al., 2019). For bleach-synchronization, gravid adults were dissolved in 20% sodium hypochlorite solution. The surviving *C. elegans* embryos were washed with M9 buffer 2 times and let hatch with gentle rocking in M9 buffer at 20°C overnight. The next day, L1 larvae were distributed onto NGM plates seeded with OP50 and grown at 20°C. Alternatively, animals were age-synchronization by egg laying; adult animals were allowed to lay eggs for 2–3 h and removed again from the plates. Embryos were grown at 20°C and assayed at the indicated days of life. Temperature sensitive strains were maintained at 15°C. All age-synchronization was done by egg laying at 15°C, embryos were then moved to the indicated temperature and allowed to develop.

### Cloning of *C. elegans* expression constructs and generation of transgenic animals

The cDNA sequence of *hsp-110* was amplified using primers with appropriate restriction sites by PCR from a L4440 plasmid (Vidal *C. elegans* ORF-RNAi library, Source BioScience) and cloned into a pCR-Blunt vector in sense and antisense direction separated by an unrelated 68 bp spacer sequence. The inverted repeat sequence was then subcloned by restriction digestion into the pPD30_38 *C. elegans* expression vector containing the *unc-54* promoter and 3’UTR sequences. The *pPD30_38_hsp-110* hairpin construct was injected together with a pharyngeal co-expression marker (*myo-2p::CFP*) into young adult N2 hermaphrodites to generate transgenic lines carrying an extrachromosomal array, which was subsequently integrated by gamma irradiation. Successfully integrated lines were backcrossed at least 5 times into the WT N2 background.

For constructing the following *C. elegans* expression constructs, the MultiSite Gateway Three-Fragment Vector Construction Kit (Thermo Fisher Scientific) was used. To generate the expression plasmids for *sse1* and *sse1^K69M^* their cDNA flanked by recombination sites suitable for gateway cloning was amplified from yeast expression plasmids pCA568 and pCA573 (Andreasson et al., 2008) by PCR. The *fes1* ORF was amplified from a pCA528 expression plasmid. The fragments were inserted into the gateway cloning entry vector pDONR221 according to manufacturer’s protocol. pDONR P4-P1R *myo-3* promoter region and pDONR P2R-P3 *unc-54* 3 UTR entry vectors were generated previously (Nussbaum-Krammer et al., 2013). The final *myo-3p::sse1/sse1^K69M^::unc-54 3’UTR* and *myo-3p::fes1::unc-54 3’UTR* were assembled by recombining the entry clones into the pDEST R4-R3 II vector (Thermo Fisher Scientific) according to the manufacturer’s protocol and sequenced after transformation. The expression plasmids (50 ng/µL) together with a pharyngeal co-expression marker *myo-2p::RFP* (Addgene) were injected into young adult α-Syn::YFP;HPI hermaphrodites to generate transgenic lines carrying an extrachromosomal array. The FLUCSM::GFP;HPI;SSE1/SSE1^K69M^ strains were generated by genetic crossing.

The endogenous *hsp-110* locus was tagged with GFP as described in Dickinson et al. (Dickinson et al., 2015). In short, the N_20_ sequence targeting the 3’ end of the *hsp-110* gene (5’-TCTATCACTTTTCGTATCAA-3’) was inserted into the Cas9–sgRNA vector pDD162 (Addgene) by site directed mutagenesis. Approx. 500 bp *hsp-110* homology arms were amplified from genomic DNA by PCR and inserted into the AvrII and SpeI double digested FP-SEC Vector pDD282 (Addgene). N2 animals were injected with a mix containing the Cas9-sgRNA vector, the *hsp-110::GFP* template, and co-injection markers, screened for positive knock-in events, before the selection cassette was removed according to the published protocol (Dickinson et al., 2015).

The quality of all constructs was validated by Sanger sequencing. Strains used in this study are listed in Table S1.

### Heat shock experiments

To induce the heat shock response (HSR), age-synchronized animals were subjected to heat stress for 3h at 33°C (+HS) in a water bath then returned to 20°C or left at 20°C (-HS). Images were taken after 2h and 24h. The -HS controls were imaged at the same time point as the +HS animals after 24h at 20°C.

For quantitative real-time (qRT)-PCR experiments, WT N2 or transgenic animals (*sid-1(pk3321)* ± HSP-110 hairpins (HP I+II)) were synchronized by bleaching and grown at 20°C until the first day of adulthood. Animals were subjected to a 1h HS at 33°C and returned to 20°C for one hour (+HS) or left at 20°C (-HS) prior to RNA extraction.

### RNA extraction and cDNA synthesis

Animals were washed off + and -HS plates with M9, transferred into 15 ml Falcon tubes and washed 3 times to remove bacteria and larvae. Worm pellets were transferred to 2 mL screw cap tubes (MP Biomedicals) and total RNA was extracted with TRIzol reagent (Invitrogen). 250 µL reagent and a similar volume of zirconia beads (0.7 mm, BioSpec Products; Carl Roth) was added to each worm pellet and mechanically lysed using a FastPrep-24 homogenizer (MP Biomedicals; 6 m/s, 45 s at 4°C). The lysate was transferred to Eppendorf tubes containing 250 µL TRIzol and 100 µL chloroform was added. The tubes were vortexed, incubated at RT for 3 min followed by a centrifugation for 15 min, 11,200 x g at 4°C. The aqueous top layer was transferred to fresh Eppendorf tubes and an equal volume of 70% ethanol was added. The containing RNA was purified using the RNeasy Mini Kit (QUIAGEN) according to the manufacturer’s protocol. RNA quality was assessed on a spectrophotometer. 1 µg RNA was cleared from gDNA by DNase I digestion (First strand cDNA Synthesis kit, Thermo Fisher Scientific). 200 ng DNase I digested RNA was used to generate cDNA with the SuperScript III First-Strand Synthesis System (Thermo Fisher Scientific) according to manufacturer’s protocol using the supplied random hexamer primers.

### Real-time PCR

qRT-PCR was performed with SYBR™ Green PCR Master Mix (Thermo Fisher Scientific), a primer concentration of 5 µM, and 2 ng cDNA as template in a 10 µL reaction volume in a LightCycler 480 II (Roche). Each primer pair condition combination was run in triplicate and the whole assay was performed 4 times. Ct values were obtained from the LightCycler 480 Software (Roche, ver. 1.5.0.39) and relative quantification of gene expression levels calculated according to Livak and Schmittgen et al. (Livak and Schmittgen, 2001) with *cdc-42* serving as the reference gene and Ct values of N2 -HS samples as calibrator samples.

### Immunostaining

Immunostaining was performed as previously described (Ben-Zvi et al., 2009; Duerr, 2013; Gidalevitz et al., 2006). Shortly, animals were freeze cracked using polylysine coated slides (Thermo Fisher Scientific) and fixed in acetone. After fixation samples were moved to filter baskets as described by (Bolkova and Lanctot, 2016). Nematodes were stained with Alexa Flour^TM^ 647-phalloidin (Thermo Fisher Scientific) and with either anti-paramyosin [5-23] or anti-myosin heavy chain A [5-6] antibodies (Ardizzi and Epstein, 1987; Ben-Zvi et al., 2009; Gidalevitz et al., 2006). Secondary antibodies were IgG-goat anti-mouse Alexa Flour 488 (Thermo Fisher Scientific). Animals were imaged as described using 488- and 640-nm for excitation.

### Fluorescence microscopy

For imaging, age-synchronized animals were mounted on 5–10% agarose (VWR Chemicals) pads and immobilized in a solution containing 2.5 mM levamisole (Applichem) and nanosphere size standards (100 nm polystyrene beads, Thermo Fisher Scientific).

High sensitivity microscopy was performed using a Leica DMi8SD AF spinning disc microscope (Leica Microsystems) equipped with 488 nm (50 mW), 561 nm (50 mW) and 640 nm lasers (100mW), an Hamamatsu Orca Flash 4.0 LT (C11440-42U) camera and the MetaMorph Advanced Acquisition software (Molecular Devices) (Figure 1A; 2G; 4A,D; S2B; S4A; S5B-E; S6B,C). Z-stacks (0.2 µm steps) and single plane images were taken using a HC PL APO 63x/1.40–0.60 oil or 100x/1.40 Oil CS2 objective. For Figures 1B, 2A, 3A, 3D, and S1A, a widefield imaging system (Xcellence IX81, Olympus, Japan) equipped with an UPlanSApo 40x/0.95 and an Apo N 60x/1.49 oil objective was employed. Z-stacks (0.2 µm steps) or single plane images were recorded with an Orca-R2 EMCCD camera (Hamamatsu, Japan) using Xcellence software (Olympus, Japan). For Figure 2D and S1D, a Leica M205 FA microscope with a Leica DFC310FX camera and the Leica DFC Twain Software were used (Leica Microsystems). All further processing of acquired images was performed with ImageJ software (NIH). If not otherwise indicated, maximum projections of z-stacks are shown.

### Quantification of foci

Heat shock-induced FLUCSM and α-Syn::YFP foci were quantified using the widefield imaging system (Xcellence IX81, Olympus) and an Apo N 60x/1.49 oil objective. Total ROIs (region of interests) were defined by manually selecting muscle cells in maximum intensity projections of deconvolved z-stacks (Wiener filter, Xcellence software). All original maximum intensity projections were processed by applying the same thresholding. For the FLUCSM quantification, the “analyze particles” tool in ImageJ (NIH) was used to determine the FLUCSM::GFP foci area relative to total ROI: Percent disaggregation was calculated as follows: 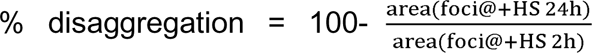. For the α-Syn::YFP foci, the “analyze particles” function was also employed to calculate the mean foci fluorescence and foci area relative to the total ROI muscle area. Relative fluorescence of foci (relFluoFoci) was defined as the product of mean fluorescence intensity of foci area and foci area size divided through total ROI size: 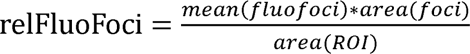. Quantification of the number of Q35 foci was performed using a Leica M205 FA stereomicroscope (Leica Microsystems). The total number of Q35::YFP foci of all muscle cells were counted manually in age-synchronized worms. Quantification of the number of FLUCSM foci during aging was performed using a Leica DMi8SD AF spinning disc microscope (Leica Microsystems). The number of animals that were positive for FLUCSM foci were counted manually.

### Worm lysis, western blot and signal quantification

Worm lysis was performed as described (Sandhof et al., 2019). In short, worms were mechanically lysed using a FastPrep-24 homogenizer (MP Biomedicals; 6.5 m/s, 60 s at 4°C) in lysis buffer buffer (20 mM Tris, pH 7.5; 10 mM ß-mercaptoethanol; 1% Triton X-100; supplemented with complete protease inhibitor cocktail (Roche)) with an appropriate amount of zirconia beads (0.7 mm, BioSpec Products; Carl Roth). The lysates were transferred into fresh Eppendorf tubes to remove beads and centrifuged (1000 x g for 2 min at 4°C) in a tabletop centrifuge to remove carcasses. The protein concentration was determined using protein assay dye reagent concentrate (Bio-Rad). Proteins were separated under denaturing conditions by SDS-PAGE and transferred onto a PVDF membrane (Carl Roth) by semi-dry or wet blotting using standard protocols. For transgene detection, mouse monoclonal anti- α-Syn antibody (211, Santa Cruz Biotechnology) or anti-GFP antibody (clone B34, Biolegend) were used. Anti-actin antibody (clone C4, Sigma-Aldrich) was used as loading control. Alkaline phosphatase (AP)-conjugated anti-mouse IgG secondary antibodies (Vector Laboratories) were used for subsequent ECF-based detection (GE Healthcare). Western blot signal intensity was analyzed using the ImageStudioLite software (version 5.2.5, Li-Cor Biosciences) on results from 3-5 independent experiments.

### Thrashing assay

In order to evaluate the physiological consequences of the knockdown of HSP-110 by expression of the hairpin construct in temperature sensitive mutants and in combination with the expression of alpha synuclein or polyglutamine, the function of muscle cells was assessed by a thrashing assay as previously described (Sandhof et al., 2019). Briefly, age-synchronized animals were transferred into M9 buffer from NGM plates and were given 1 min to adapt to the new environment. The number of swimming movements (thrashes) within 30 s was counted to determine the swimming speed. One thrash was defined as the entire motion from one side to the other. 10–15 worms were examined for each strain in 3 biological replicates.

### Statistical analysis

For all experiments, at least 3 biological replicates were obtained for each strain tested. GraphPad Prism software was used to create graphs and to analyze the data. Data are presented as mean ± standard error of the mean (SEM). To determine whether there are statistically significant differences between strains or treatments, the following statistical tests were performed. The two-way analysis of variance (ANOVA) with Bonferroni posttests, Dunnett’s or Tukey’s multiple comparisons test were used to compare 2 variables (strains and age) with each another. The one-way ANOVA together with a Dunnett, or Tukey’s multiple comparisons test was used to compare every mean to a control mean.

