## Supplemental Figure Legends for "HSP110 dependent HSP70 disaggregation machinery mediates prion-like propagation of amyloidogenic proteins in metazoa"

**Supplemental Material**

**Figure S1. The *sid-1(pk3321)* loss-of-function allele does not reduce α-Syn and Q35 foci formation and toxicity**

**A** Maximum intensity projections of fluorescent microscopy z-stacks of 5-day-old animals expressing α-Syn::YFP in the WT and *pk3321* mutant background. Scale bars: 10 µm.

**B** Quantification of α-Syn::YFP foci in 4-, 5-, and 6-day-old worms. RelFluoFoci is displayed (mean ± SEM in %). Statistical analysis was done using 1-way ANOVA and post-hoc test with Tukey adjustment for multiple comparisons. n.s. = not significant, ** $p \leq 0.01$.

**C** Quantification of motility as a measure for aggregate toxicity. Displayed is the mean amount of body bends per 30 seconds ± SEM. Statistical analysis was done using two-way ANOVA with Bonferroni's adjustment for multiple comparisons. n.s. = not significant, ** $p \leq 0.01$.

**D** Maximum intensity projections of fluorescent microscopy z-stacks of 6-day-old animals expressing Q35::YFP in the WT and *pk3321* mutant background. Scale bars: 100 µm.

**E** Manual count of Q35::YFP foci in the WT and *pk3321* mutant background (mean ± SEM). Statistical analysis was done using two-way ANOVA with Dunnett's multiple comparison test. n.s. = not significant.

**F** Quantification of motility as a measure for aggregate toxicity. Displayed is the mean number of body bends per 30 seconds ± SEM. Statistical analysis was done using one-way ANOVA with Tukey's multiple comparison test. n.s. = not significant.

**Figure S2. KD of Hsp110 does not result in a compensatory activation of the heat shock response**

**A** Quantification of mRNA levels in 4-day-old animals expressing the indicated transgenes or harboring the indicated mutation relative to the -HS control of major HS-inducible genes (2 HSP70s, *C12C8.1* and *F44E5.4*, and the small HSP *hsp-16.2*) by real-time (RT)-PCR (mean ± SEM). Statistical analysis was done using two-way ANOVA with Bonferroni's multiple comparison test. n.s. = not significant.

**B** Experimental set-up: Age-synchronized 4-day-old animals were subjected to heat stress for 3h at 33°C (+HS) then returned to 20°C or left at 20°C (-HS), respectively. Arrows indicate imaging time points. The -HS controls were imaged at the same time point as the +HS animals after 2h at 20°C. Fluorescent microscopy images of animals expressing GFP under the HS-inducible *hsp-16.2* promoter in the control and HSP-110 KD background. Scale bars: 25 µm. The KD of HSP-110 did not evoke an heat shock response under ambient growth conditions (only autofluorescence of the gut granules (*) is visible in -HS strains), but animals were still able to mount an HSR under acute heat stress (+HS). All strains harbor the *sid-1(pk3321)* allele.

**Figure S3. Total α-Syn and Q35 protein levels do not change upon KD of HSP-110**

**A, B** and **C** Total lysates of age-synchronized strains (**A**: day 5; **B**: day 5 **C**: day 6) expressing the indicated transgenes. α-Syn::YFP or Q35::YFP were detected with an anti-GFP antibody and anti-α-actin antibody was used as loading control. *cross-reactivity with the myo-2p::CFP co-injection marker expressed in the hairpin animals. Respective bar graphs display protein levels relative to the control (mean ± SEM in %). Statistical analysis was done using one-way ANOVA with Tukey's multiple comparison test. n.s. = not significant. The KD of HSP-110 did not significantly affect transgene levels. All strains harbor the *sid-1(pk3321)* allele.

**Figure S4. HSP-110 HP expression impairs cellular protein folding homeostasis**

**A** Maximum intensity projections of confocal z-stacks of 4-day-old animals expressing the indicated transgenes or harboring the indicated temperature-sensitive (ts) mutations. Muscle cells were stained with anti-paramyosin (green), and Alexa Flour 647-phalloidin (purple). Scale bar: 5 µm. Paramyosin misfolding revealed that co-expression of the *hsp-110* hairpins exposed the ts mutant phenotype of paramyosin(ts) [*unc-15*(*e1402*)] at the permissive temperate of 15°C.

**B** and **C** Quantification of motility as a measure for toxicity. Displayed is the mean ± SEM of body bends/30s of animals expressing the indicated transgenes or harboring the indicated mutations. Statistical analysis was done using one-way ANOVA with Dunnett's multiple comparison test. ** $p \leq 0.01$, *** $p \leq 0.001$. Co-expression of the *hsp-110* hairpins exposed the ts mutant phenotypes already at the permissive temperate of 15°C, resulting in a significant increase in movement defects. All strains harbor the *sid-1(pk3321)* allele.

**Figure S5. HSP-110 HP expression does not affect WT myosin or paramyosin folding**

**A** Quantification of motility as a measure for transgene toxicity. Displayed is the mean ± SEM of body bends/30s of animals expressing the indicated transgenes or harboring the indicated mutations. Statistical analysis was done using one-way ANOVA with Dunnett's multiple comparison test. Animals that express the *hsp-110* hairpins show normal movement at 15°C and 25°C.

**B, C, D** and **E** Maximum intensity projections of confocal z-stacks of 4-day-old animals expressing the indicated transgenes or harboring the indicated mutations. Muscle cells were stained with **B** and **C** anti-myosin (green) or **D** and **E** anti-paramyosin (green), and Alexa Flour 647-phalloidin (purple). Scale bar: 5 μm. Animals expressing the *hsp-110* hairpins exhibited a normal myosin and paramyosin structure at 15°C and 25°C. All strains harbor the *sid-1(pk3321)* allele.

**Figure S6. HSP-110 HP expression impairs cellular protein folding homeostasis during aging**

**A** Quantification of motility as a measure for transgene toxicity. Displayed is the mean ± SEM amount of body bends/30s of animals expressing the indicated transgenes or harboring the indicated mutations. Statistical analysis was done using two-way ANOVA with Tukey's multiple comparison test. * $p \leq 0.05$, ** $p \leq 0.01$. Expression of the hsp-110 hairpins causes movement defects with increasing age at 25°C.

**B** Single plane confocal fluorescent microscopy images of 10-day-old animals harboring the indicated transgenes or endogenously tagged proteins. Muscle cells are outlined. Muscle cell specific HSP-110::GFP depletion persisted during aging in HP animals. Scale bars: 10 μm.

**C** Maximum intensity projections of confocal z-stacks of 12-day-old animals expressing the indicated transgenes or mutations. Scale bar: 10 μm. α-Syn::YFP foci number remained low and did not increase in old HPI and HPII animals.

**D** Quantification of the ratio of α-Syn::YFP foci in 12-day-old animals to 5-day-old animals. The ratio of the product of mean foci fluorescence and foci area relative to muscle area from 12-day-old to 5-day-old animals is displayed (relFluoFoci). Data are shown as mean ± SEM. All strains harbor the *sid-1(pk3321)* allele.
