## Supplementary figures and images for "HSP110 dependent HSP70 disaggregation machinery mediates prion-like propagation of amyloidogenic proteins in metazoa"

### Figure S1

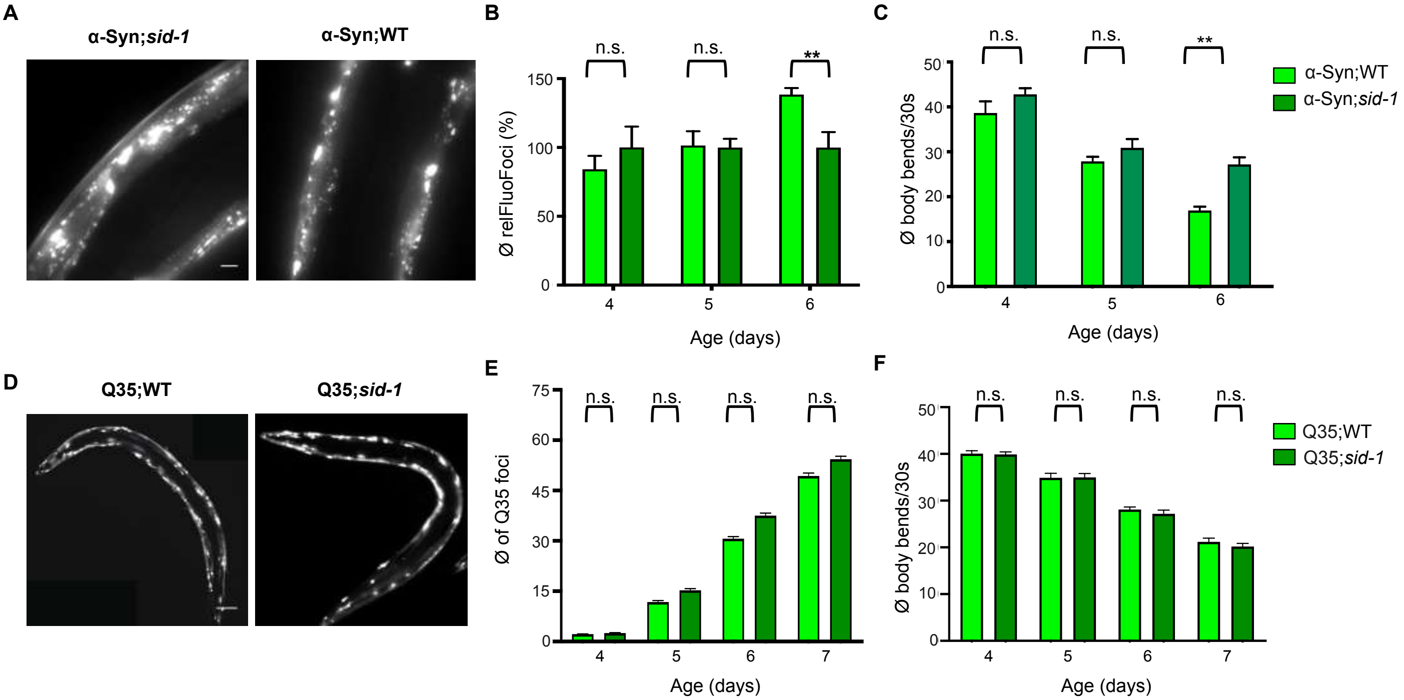

### Figure S2

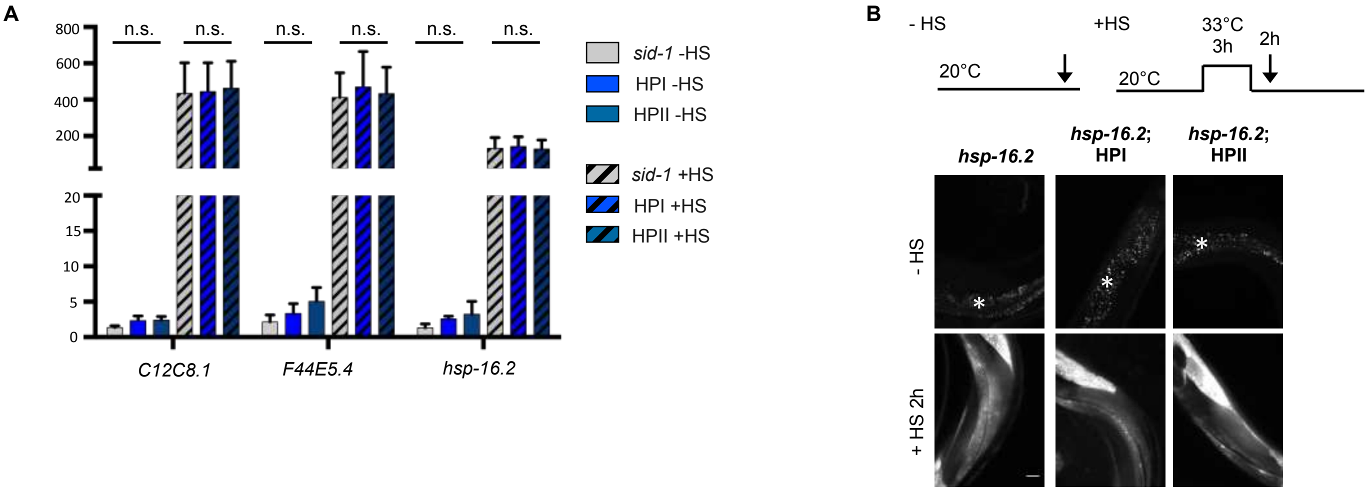

### Figure S3

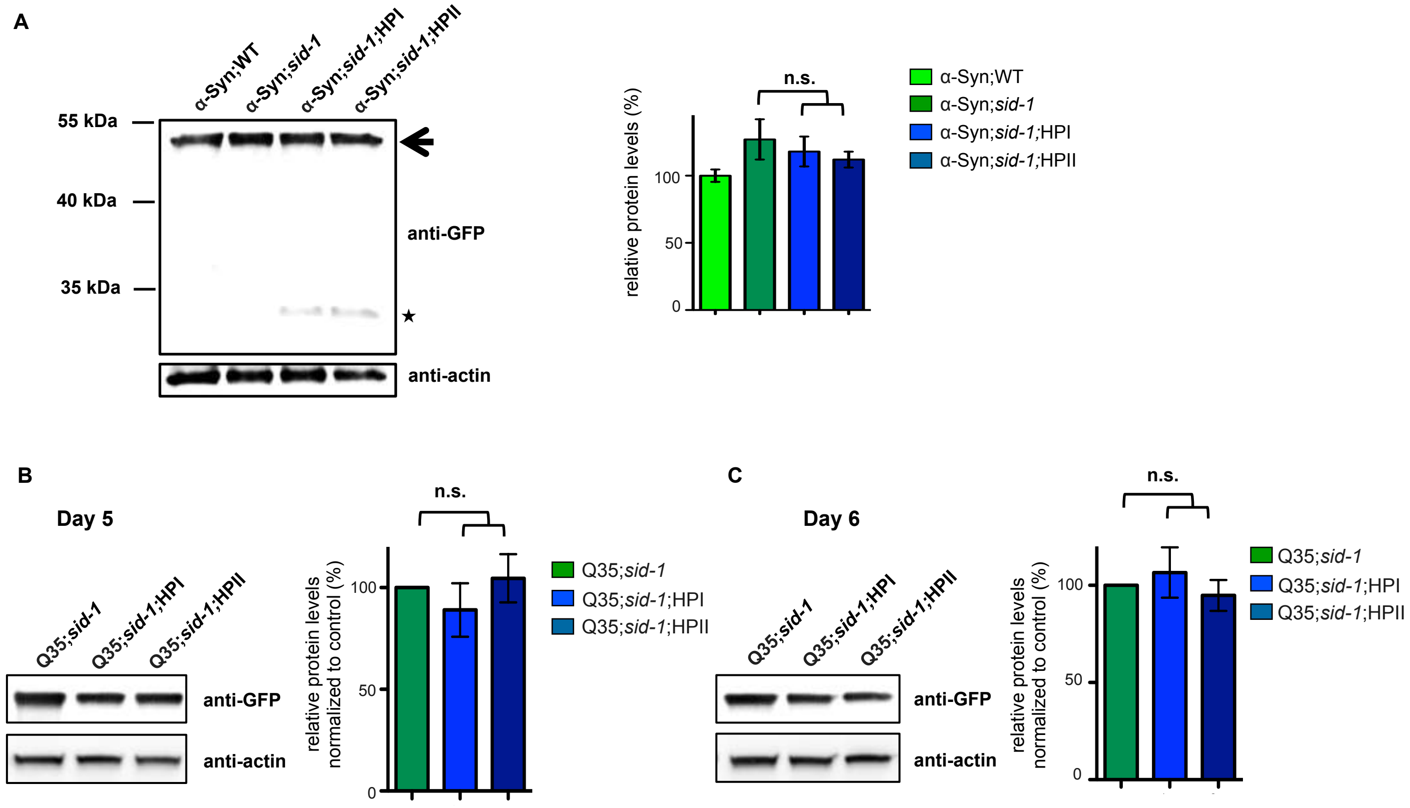

### Figure S4

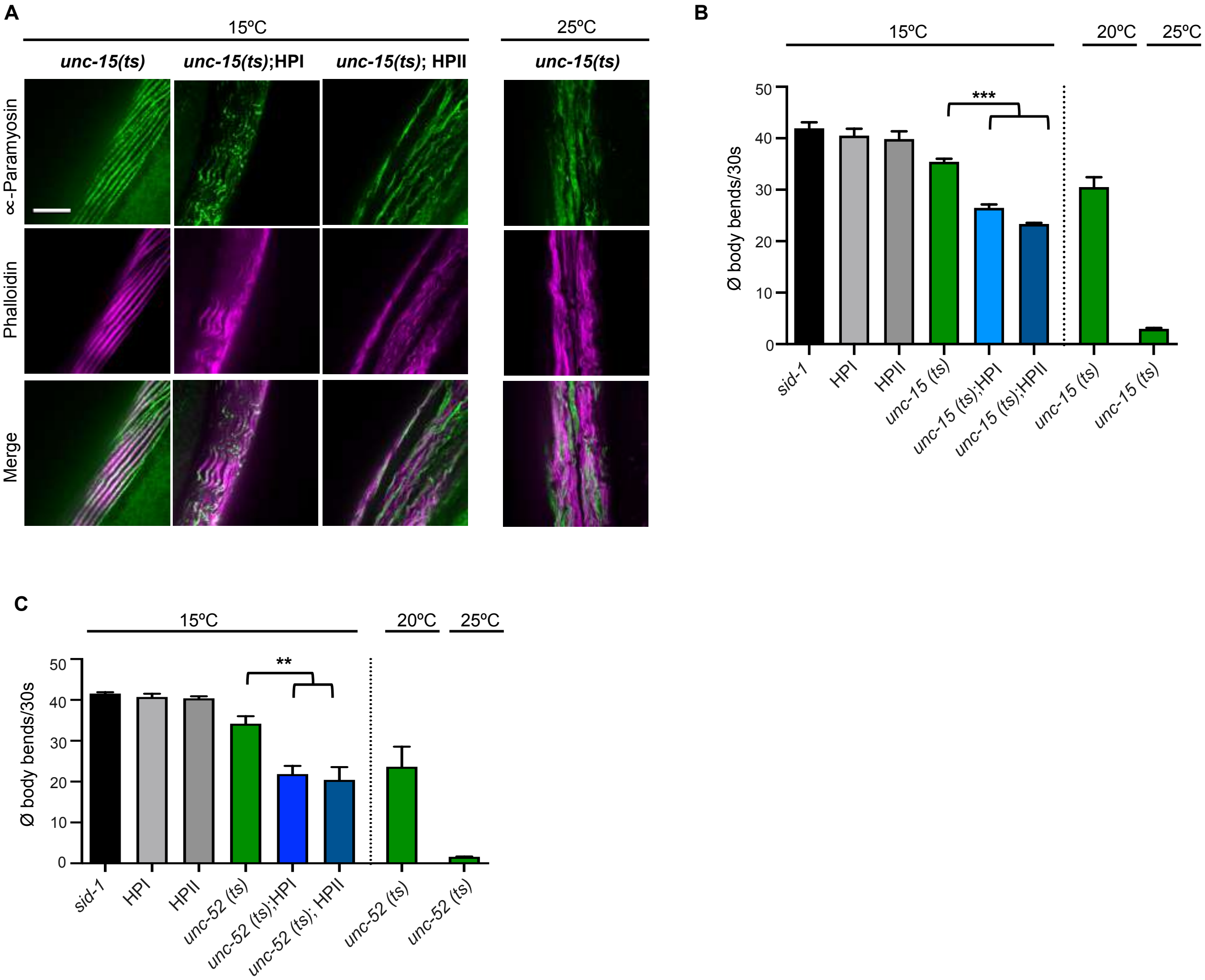

### Figure S5

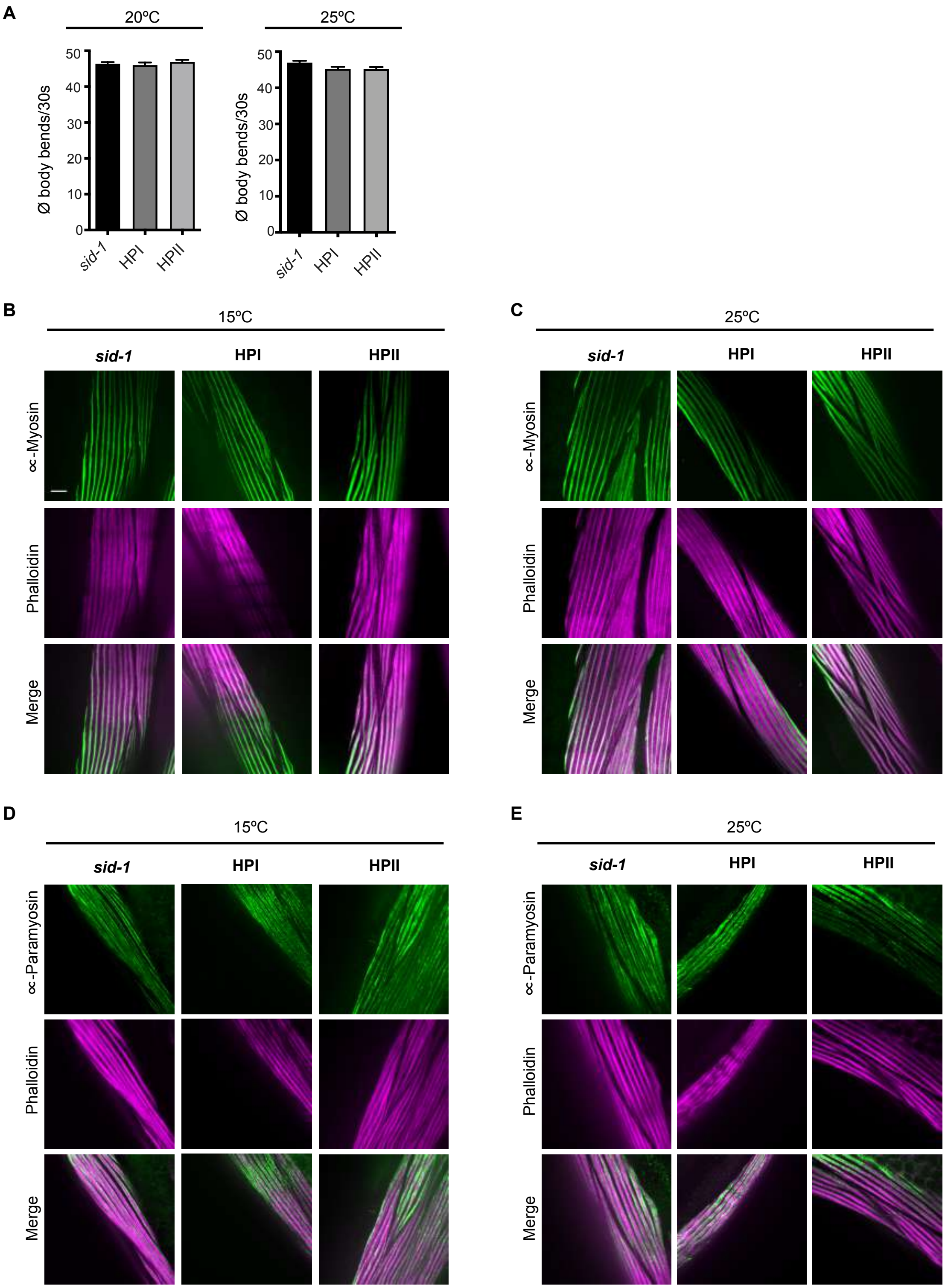

### Figure S6

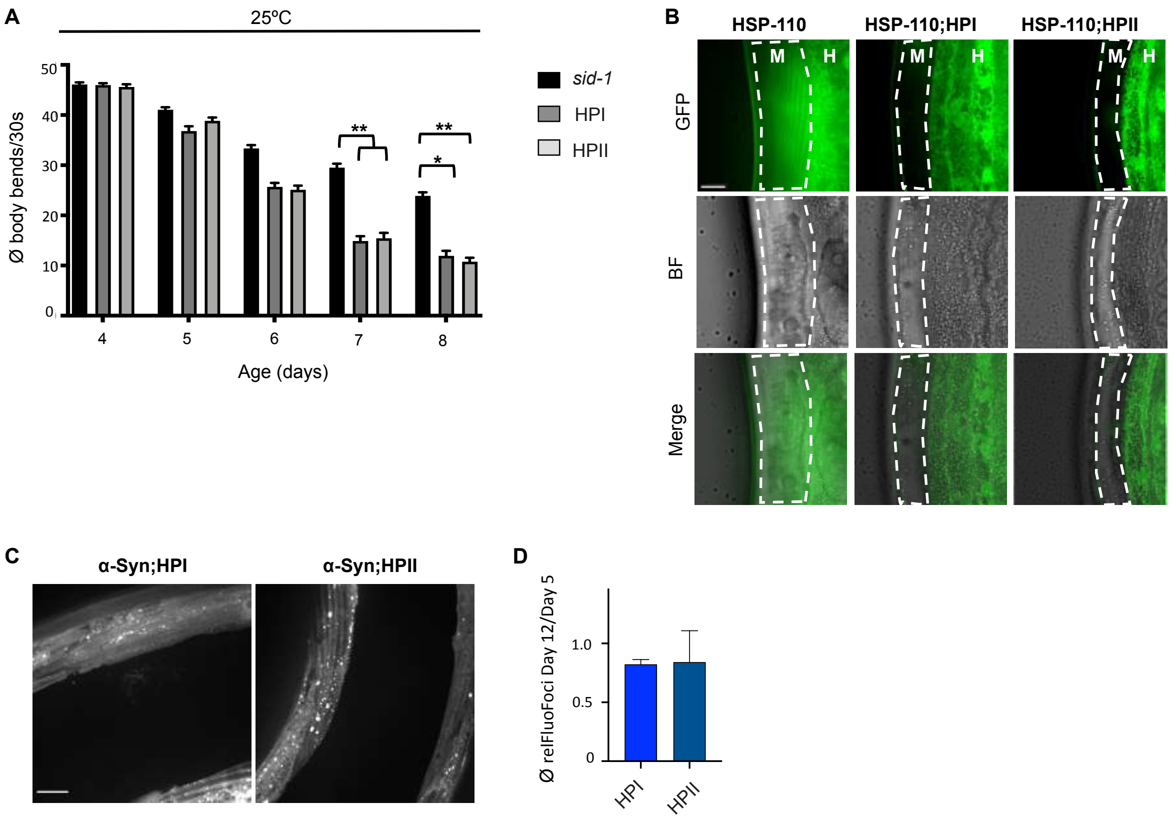
