## Supplementary material for "HSP110 dependent HSP70 disaggregation machinery mediates prion-like propagation of amyloidogenic proteins in metazoa": Table S1

| Strain name | Genotype | Source | Reference |
|---|---|---|---|
| AM135 | rmIs127[unc-54p::Q0::YFP] | Dr. Rick Morimoto | Morley et al. (2002),PNAS |
| AM137 | rmIs129[unc-54p::Q24::YFP] | Dr. Rick Morimoto | Morley et al. (2002),PNAS |
| AM140 | rmIs132[unc-54p::Q35::YFP] I. | Dr. Rick Morimoto | Morley et al. (2002),PNAS |
| OW40 | zgIs15[P(unc-54)::asyn:: YFP]IV | Dr. Rick Morimoto | Van Ham et al. (2008), PLoS Genet. |
| AM1218 | rmIs400[myo-3p::alpha-synuclein WT::RFP::unc-54 3'UTR] | Nussbaum-Krammer lab | Sandhof et al. (2019), Autophagy |
| NL3321 | sid-1[pk3321 V] | CGC | |
| CL2070 | dvIs70 [hsp-16.2p::GFP + rol-6(su1006)]. | CGC | |
| CB1301 | unc-54(e1301) I. | Dr. Rick Morimoto | |
| CB1402 | unc-15(e1402) I. | Dr. Rick Morimoto | |
| HE250 | unc-52(e669su250) II. | Dr. Rick Morimoto | |
| FUH134 | marIs134 [unc-54p::luciferase (R188Q)::gfp + rol-6(su1006)] | Dr. Rick Morimoto | |
| CNK300 | prpIs61 [unc-54p::hsp110-HP I-4 + myo-2p::CFP] + sid-1[pk3321 V] | This study | This study |
| CNK301 | prpIs62 [unc-54p::hsp110-HP III-6 + myo-2p::CFP] + sid-1[pk3321 V] | This study | This study |
| CNK302 | prpIs63[Hsp110::GFP^3xFlag] + sid-1 [pk3321 V] | This study | This study |
| CNK303 | prpIs63[Hsp110::GFP^3xFlag] + prpIs61 [unc-54p::hsp110-HP I-4 + myo-2p::CFP] + sid-1[pk3321 V] | This study | This study |
| CNK304 | prpIs63[Hsp110::GFP^3xFlag] + prpIs62 [unc-54p::hsp110-HP III-6 + myo-2p::CFP] + sid-1[pk3321 V] | This study | This study |
| CNK305 | marIs134 [unc-54p::luciferase (R188Q)::gfp + rol-6] + sid-1[pk3321 V] | This study | This study |
| CNK306 | marIs134 [unc-54p::luciferase (R188Q)::gfp + rol-6] + prpIs61 [unc-54p::hsp110-HP I-4 + myo-2p::CFP] + sid-1[pk3321 V] | This study | This study |
| CNK307 | marIs134 [unc-54p::luciferase (R188Q)::gfp + rol-6] + prpIs62 [unc-54p::hsp110-HP III-6 + myo-2p::CFP] + sid-1[pk3321 V] | This study | This study |
| CNK308 | zgIs15[P(unc-54)::asyn:: YFP]IV + sid-1[pk3321 V] | This study | This study |
| CNK309 | zgIs15[P(unc-54)::asyn:: YFP]IV + prpIs61 [unc-54p::hsp110-HP I-4 + myo-2p::CFP] + sid-1[pk3321 V] | This study | This study |

| | | | |
|---|---|---|---|
| CNK310 | zgIs15[P(unc-54)::asyn:: YFP]IV + prpIs62 [unc-54p::hsp110-HP III-6 + myo-2p::CFP] + sid-1[pk3321 V] | This study | This study |
| CNK311 | rmIs132[unc-54p::Q35::YFP] I + sid-1[pk3321 V] | This study | This study |
| CNK312 | rmIs132[unc-54p::Q35::YFP] I + prpIs61 [unc-54p::hsp110-HP I-4 + myo-2p::CFP] + sid-1[pk3321 V] | This study | This study |
| CNK313 | rmIs132[unc-54p::Q35::YFP] I + prpIs62 [unc-54p::hsp110-HP III-6 + myo-2p::CFP] + sid-1[pk3321 V] | This study | This study |
| CNK20 | rmIs400[myo-3p::alpha-synuclein WT::RFP::unc-54 3'UTR] + sid-1[pk3321 V] | This study | This study |
| CNK31 | rmIs400[myo-3p::alpha-synuclein WT::RFP::unc-54 3'UTR] + prpIs61 [unc-54p::hsp110-HP I-4 + myo-2p::CFP] + sid-1[pk3321 V] | This study | This study |
| CNK24 | rmIs400[myo-3p::alpha-synuclein WT::RFP::unc-54 3'UTR] + prpIs62 [unc-54p::hsp110-HP III-6 + myo-2p::CFP] + sid-1[pk3321 V] | This study | This study |
| CNK314 | dvIs70[myo-3::Abeta 1-42 wt::3' UTR(long) + mtl-2::GFP; hsp16.2p::GFP; rol-6(su1006)]+ sid-1[pk3321 V] | This study | This study |
| CNK315 | dvIs70[myo-3::Abeta 1-42 wt::3' UTR(long) + mtl-2::GFP; hsp16.2p::GFP; rol-6(su1006)]+ prpIs61 [unc-54p::hsp110-HP I-4 + myo-2p::CFP] + sid-1[pk3321 V] | This study | This study |
| CNK316 | dvIs70[myo-3::Abeta 1-42 wt::3' UTR(long) + mtl-2::GFP; hsp16.2p::GFP; rol-6(su1006)]+ prpIs62 [unc-54p::hsp110-HP III-6 + myo-2p::CFP] + sid-1[pk3321 V] | This study | This study |
| CNK317 | prpEx49 [myo-3p::Sse1::unc-54 3'UTR + myo-2p::mCherry]; zgIs15[P(unc-54)::asyn:: YFP]IV + prpIs61 [unc-54p::hsp110-HP I-4 + myo-2p::CFP] + sid-1[pk3321 V] | This study | This study |
| CNK318 | prpEx52 [myo-3p::Sse1 K69M::unc-54 3'UTR + myo-2p::mCherry]; zgIs15[P(unc-54)::asyn:: YFP]IV + prpIs61 [unc-54p::hsp110-HP I-4 + myo-2p::CFP] + sid-1[pk3321 V] | This study | This study |
| CNK319 | prpEx53 [myo-3p::Fes1::unc-54 3'UTR + myo-2p::mCherry]; zgIs15[P(unc-54)::asyn:: YFP]IV + prpIs61 [unc-54p::hsp110-HP I-4 + myo-2p::CFP] + sid-1[pk3321 V] | This study | This study |

| | | | |
|---|---|---|---|
| CNK320 | prpEx49 [myo-3p::Sse1::unc-54 3'UTR + myo-2p::mCherry]; marIs134 [unc-54p::luciferase$^{R188Q}$:: eGFP + rol-6] + prpIs61 [unc-54p::hsp110-HP I-4 + myo-2p::CFP]+ sid-1[pk3321 V] | This study | This study |
| CNK321 | prpEx52 [myo-3p::Sse1$^{K69M}$::unc-54 3'UTR + myo-2p::mCherry]; marIs134 [unc-54p::luciferase$^{R188Q}$:: eGFP + rol-6] + Is[unc-54p::hsp110-HPI-4 + myo-2p::CFP] + sid-1[pk3321 V] | This study | This study |
| CNK322 | unc-54(e1301) I. + sid-1[pk3321 V] | This study | This study |
| CNK131 | prpIs61 [unc-54p::hsp110-HP I-4 + myo-2p::CFP] + sid-1[pk3321 V] + unc-54(e1301) I | This study | This study |
| CNK130 | prpIs62 [unc-54p::hsp110-HP III-6 + myo-2p::CFP] + sid-1 [pk3321] + unc-54(e1301) I | This study | This study |
| CNK323 | unc-15(e1402) I + sid-1[pk3321 V] | This study | This study |
| CNK132 | prpIs61 [unc-54p::hsp110-HP I-4 + myo-2p::CFP] + sid-1 [pk3321] + unc-15(e1402) I | This study | This study |
| CNK133 | prpIs62 [unc-54p::hsp110-HP III-6 + myo-2p::CFP] + sid-1 [pk3321] + unc-15(e1402) I | This study | This study |
| CNK324 | unc-52(e669su250) II + sid-1[pk3321 V] | This study | This study |
| CNK134 | prpIs61 [unc-54p::hsp110-HP I-4 + myo-2p::CFP] + sid-1[pk3321 V] + unc-52(e669su250) I | This study | This study |
| CNK135 | prpIs62 [unc-54p::hsp110-HP III-6 + myo-2p::CFP] + sid-1 [pk3321] + unc-52(e669su250) I | This study | This study |
| CNK191 | prpEx49 [myo-3p::Sse1::unc-54 3'UTR + myo-2p::mCherry]; rmIs400[myo-3p::alpha-synuclein WT::RFP::unc-54 3'UTR]; prpIs61 [unc-54p::hsp110-HP I-4 + myo-2p::CFP] + sid-1[pk3321 V] | This study | This study |
| CNK192 | prpEx52 [myo-3p::Sse1 K69M::unc-54 3'UTR + myo-2p::mCherry]; rmIs400[myo-3p::alpha-synuclein WT::RFP::unc-54 3'UTR] + prpIs61 [unc-54p::hsp110-HP I-4 + myo-2p::CFP] + sid-1[pk3321 V] | This study | This study |
